# Spatial transcriptomics reveals ovarian cancer subclones with distinct tumour microenvironments

**DOI:** 10.1101/2022.08.29.505206

**Authors:** Elena Denisenko, Leanne de Kock, Adeline Tan, Aaron B. Beasley, Maria Beilin, Matthew E. Jones, Rui Hou, Dáithí Ó Muirí, Sanela Bilic, G. Raj K. A. Mohan, Stuart Salfinger, Simon Fox, Khaing Hmon, Yen Yeow, Elin S. Gray, Paul A. Cohen, Yu Yu, Alistair R. R. Forrest

## Abstract

High-grade serous ovarian carcinoma (HGSOC) is characterised by recurrence, chemotherapy resistance and overall poor prognosis. Genetic heterogeneity of tumour cells and the microenvironment of the tumour have been hypothesised as key determinants of treatment resistance and relapse. Here, using a combination of spatial and single cell transcriptomics (10x Visium and Chromium platforms), we examine tumour genetic heterogeneity and infiltrating populations of HGSOC samples from eight patients with variable response to neoadjuvant chemotherapy. By inferring gross copy number alterations (CNAs), we identified distinct tumour subclones co-existing within individual tumour sections. These tumour subclones have unique CNA profiles and spatial locations within each tumour section, which were further validated by ultra-low-pass whole genome sequencing. Differential expression analysis between subclones within the same section identified both tumour cell intrinsic expression differences and markers indicative of different infiltrating cell populations. The gene sets differentially expressed between subclones were significantly enriched for genes encoding plasma membrane and secreted proteins, indicative of subclone-specific microenvironments. Furthermore, we identified tumour derived ligands with variable expression levels between subclones that correlated or anticorrelated with various non-malignant cell infiltration patterns. We highlight several of these that are potentially direct tumour-stroma/immune cell relationships as the non-malignant cell type expresses a cognate receptor for the tumour derived ligand. These include predictions of *CXCL10-CXCR3* mediated recruitment of T and B cells to associate with the subclones of one patient and *CD47-SIRPA* mediated exclusion of macrophages from association with subclones of another. Finally, we show that published HGSOC molecular subtype signatures associated with prognosis are heterogeneously expressed across tumour sections and that areas containing different tumour subclones with different infiltration patterns can match different subtypes. Our study highlights the high degree of intratumoural subclonal and infiltrative heterogeneity in HGSOC which will be critical to better understand resistance and relapse.

## Introduction

Ovarian cancer is the eighth leading cause of cancer deaths in women worldwide ^1^. High-grade serous ovarian carcinoma (HGSOC) is the most common and lethal histologic subtype, accounting for 70-80% of ovarian cancer deaths ^2^. HGSOC is thought to be derived from both fallopian tube and ovarian surface epithelium ^3,4^ and genomically, is characterised by almost universal *TP53* mutations and copy number alterations (CNAs) ^5–8^. Notably, although several chromosomal regions are recurrently altered ^5^, and multiple genes (*FAT3, CSMD3, BRCA1, BRCA2, NF1, CDK12, GABRA6, RB1, NF1, PTEN* and *RAD51B*) are recurrently disrupted, HGSOC genomes are highly heterogeneous with most of the above alterations only found in a small fraction of tumours ^5,9–12^. Also, due to a high degree of chromosomal instability ^13^, most HGSOCs are polyclonal ^14,15^. As the cancer progresses and metastasises, clonal diversity increases which is associated with worse prognosis and development of chemoresistance ^8,9,13,16,17^.

In addition to intratumoural clonal heterogeneity, HGSOC tumours contain a diverse range of non-malignant cell types. Recently, several single-cell RNA-sequencing (scRNA-seq) studies of primary and metastatic tumours have described the cell types that make up the HGSOC tumour microenvironment ^18–23^. With these single cell profiles it is now apparent that previously reported transcriptional subtypes of HGSOC based on bulk expression measurements (mesenchymal (C1.MES), immunoreactive (C2.IMM), differentiated (C4.DIF), and proliferative (C5.PRO)) which are associated with differences in prognosis ^24^ largely reflect the degree of immune cell infiltration and the abundance of fibroblasts ^19^, rather than inherent differences in tumour cells. To determine how these non-malignant cell types might influence tumour growth and prognosis, several groups have predicted ligand-receptor interactions between stromal, immune and tumour cell populations ^23,25^. Lastly, copy number alterations can be inferred from scRNA-seq data and this strategy has been used to identify CNAs in HGSOC tumour cells ^19,21^ and reveal subclones with different CNAs ^25^.

Here we have used spatial transcriptomics (10x Genomics Visium) of HGSOC tumours to reveal the relationship between tumour subclonal genotypes and infiltration patterns by non-malignant cell types. Using CNA inference, we predict multiple regionally distinct subclones within small tumour sections (< 6.5mm^2^) and show that they often have different patterns of infiltration that correspond to previously described molecular subtypes. By integration with scRNA-seq data, we identify tumour cell derived ligands that are differentially expressed between subclones and correlated or anticorrelated with the degree of infiltration by non-malignant cell types expressing cognate receptors for the ligands. This provides a likely link between subclonal genotype differences and differential infiltration patterns.

## Results

### Spatial gene expression of HGSOC tumours

Visium spatial transcriptomics technology uses a grid of ~5,000 55μm spots containing uniquely barcoded oligo-dT primers for cDNA synthesis placed 100μm from each other to spatially sample RNAs from an overlaid tissue section. Here, we used Visium technology on sections of primary tumours collected during interval debulking surgery from eight HGSOC patients to explore their cellular composition and tumour microenvironments. The tumour samples were collected from HGSOC patients who underwent taxane- and platinum-based neoadjuvant chemotherapy (**Supplementary Figure S1**) and included three patients with poor chemotherapy response score (CRS1: patients 1, 7 & 8), three with good response (CRS3: patients 2, 3 & 5) and two with a partial response (CRS2: patients 4 & 6) ^26^. Across the eight patient samples, the number of Visium spots yielding data ranged from 1,501 to 3,584 per section, with a total of 19,990 spots detected in the dataset and a median of 2,459 genes (5,882 unique molecular identifiers, UMIs) detected per spot (**Supplementary Table ST1**). Gene expression clustering of the spots in each section identified eight to ten clusters per sample. Strikingly, the spatial distribution of the identified clusters largely mirrored morphologically distinct regions of the sections seen after haematoxylin and eosin (H&E) staining (**Figure 1a-b**). For instance, the clusters shown in orange and yellow for patient 1 in **Figure 1b** correspond to areas at the top and bottom of the tissue section that are visually distinct from the rest of the section on the H&E staining image (**Figure 1a**). This indicates that underlying these different morphologies are different proportions of cell types and states with different gene expression patterns.

**Figure 1.**
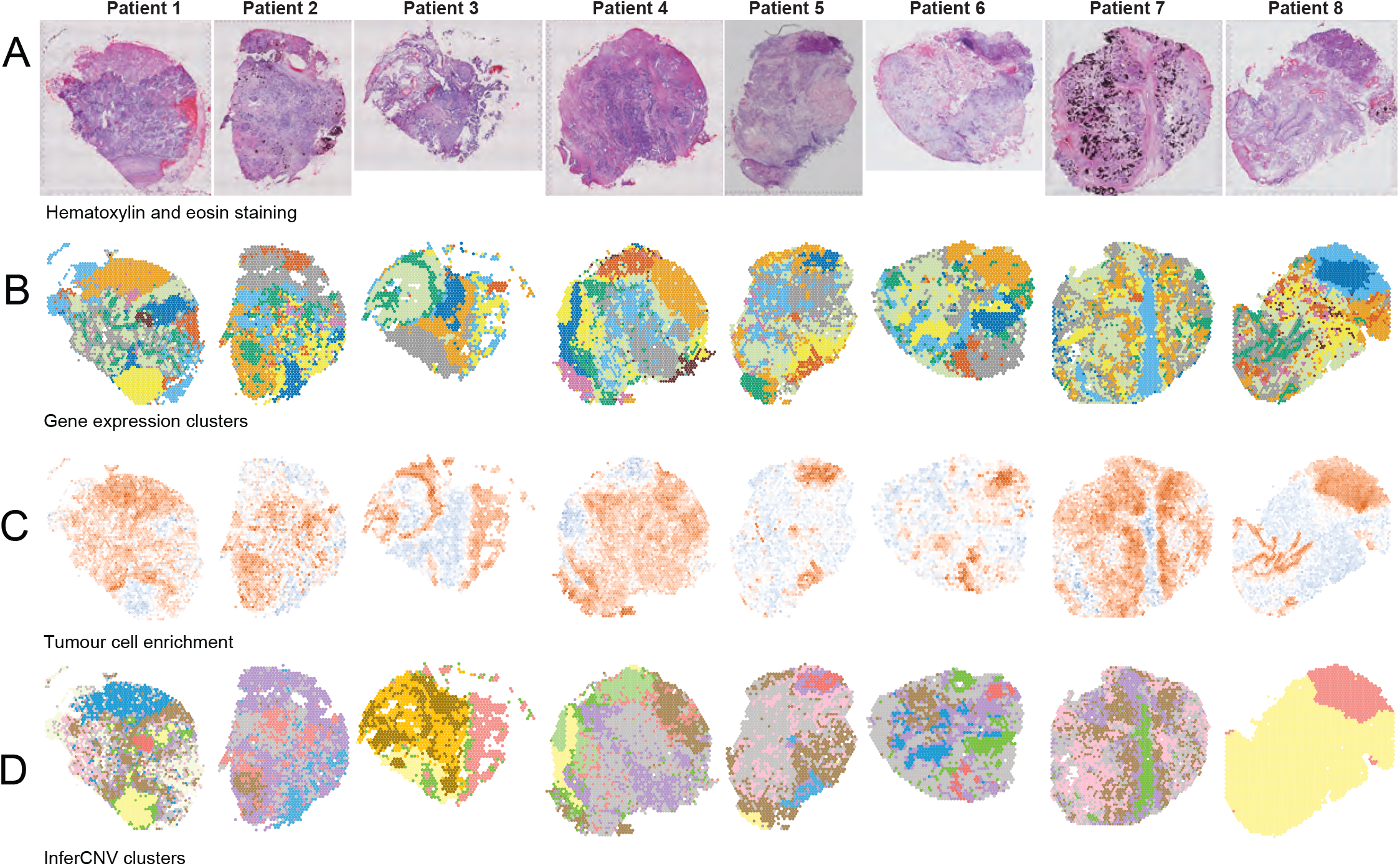
Visium profiles of eight HGSOC samples. Shown are **a**) hematoxylin and eosin stained tissue sections, **b**) gene expression-based clusters mapped onto the tissue sections, **c**) tumour cell enrichment scores calculated using Giotto ^29^. Red indicates enrichment, blue depletion, **d**) InferCNV clusters. Yellow spots in patients 1, 3, 4, 5, 8 correspond to non-malignant regions, green corresponds to border regions, other colours correspond to putative tumour subclones.

To map the location of individual cell types across each section and annotate regions as tumour or non-tumour, we performed single-cell RNA sequencing (scRNA-seq) on five additional post neoadjuvant chemotherapy HGSOC samples (**Supplementary Figure S1**). The scRNA-seq data were processed as described in **Methods** and integrated into one dataset (17,192 cells in total, with a median of 775 genes and 2,002 UMIs per cell) using Seurat ^27^. Cells were assigned seven top-level cell type annotations using scMatch ^28^ with a previously published ovarian cancer single cell dataset as a reference ^21^, followed by sub-clustering for 12 fine-grain cell types (**Methods, Supplementary Figures S2-3, Supplementary Table ST2**). From this, a tumour cell cluster and eleven non-malignant cell types including endothelial cells, B/Plasma cells, T cells, macrophages, mesothelial cells, myofibroblasts, and five fibroblast populations fibro1 (*EIF4A3, STAR*), fibro2 (*RBP1, DCN*), fibro3 (*RAMP1, CFD*), fibro4 (*CCL2*), fibro5 (*FN1, COL3A1*) were identified. We next extracted gene lists specific for each of the 12 cell types and used Giotto ^29^ to examine their spatial distribution across each Visium section (**Methods**). We also used three manually curated gene lists for neutrophils, mast cells and adipocytes as these cell types have been reported to be present in HGSOC ^30–32^ but were not observed in our scRNA-seq data. This analysis revealed distinct areas in each section with strong enrichment or depletion of tumour cells that likely correspond to the malignant and non-malignant areas of the section (**Figure 1c**). Notably, correlations between the cell type specific Giotto scores indicated co-localisation of some cell types (e.g., fibroblast, endothelial cell and myofibroblast scores correlated strongly with each other) and a strong anticorrelation between tumour cell scores and mast cells, adipocyte, B/plasma cell, neutrophil, fibroblast, myofibroblast and endothelial cell scores (**Supplementary Figure S4**). Only the mesothelial cells score had a positive correlation with the tumour cell scores, while macrophages and T cells appeared to be equally distributed in tumour and non-tumour regions (**Supplementary Figure S4**). Amongst the non-malignant cell types, endothelial cells, myofibroblasts, fibroblasts (fibro5), macrophages and B/Plasma cells were commonly enriched in the sections and their spatial patterns indicated strong within-sample heterogeneity (**Supplementary Figure S5**). In a supplementary analysis we noted that genes more highly expressed in the malignant areas of the three good response samples were predominantly expressed in B cells, T cells, macrophages, and fibro5, whereas the genes overexpressed in the malignant areas of the three poor response samples were more highly expressed in tumour cells or myofibroblasts (**Supplementary Note 1**). This is in agreement with previous antibody studies correlating B and T cell infiltration with improved survival ^33,34^. In summary, the Visium data clearly identified morphologically distinct areas of each tumour with gene expression indicative of differences in tumour cell content and infiltrating cell populations.

### Tumour subclones with unique CNAs and spatial locations

Copy number alterations (CNAs) are ubiquitous in HGSOC^5^, therefore we applied inferCNV, which detects differences in average relative expression levels for a sliding window of 101 genes ^35^, to predict CNAs for each spot and cluster spots by similar CNA profiles. Analysis of the Visium data from patient 1 predicted multiple large alterations including amplifications of parts of chromosomes 8, 12, and 20 and deletions of parts of chromosomes 6 and 19 (**Figure 2**). Clustering of spots by their inferCNV profiles identified nine clusters, of which seven appear to be malignant, one non-malignant and one with a weaker CNV signal indicative of a mixture of tumour cells and a high proportion of non-malignant cells (**Figure 2b**). Projection of the inferCNV clusters back onto the Visium slide (**Figure 2a**) revealed clear spatial separation of non-malignant (yellow), and malignant inferCNV clusters; it also revealed that the potential mixed cluster with weaker CNA signal (green) was localised to the border between the malignant and non-malignant regions. The inferCNV-based prediction of malignant and non-malignant areas was in agreement with the Giotto tumour cell enrichment score distribution (**Figure 1c**).

**Figure 2.**
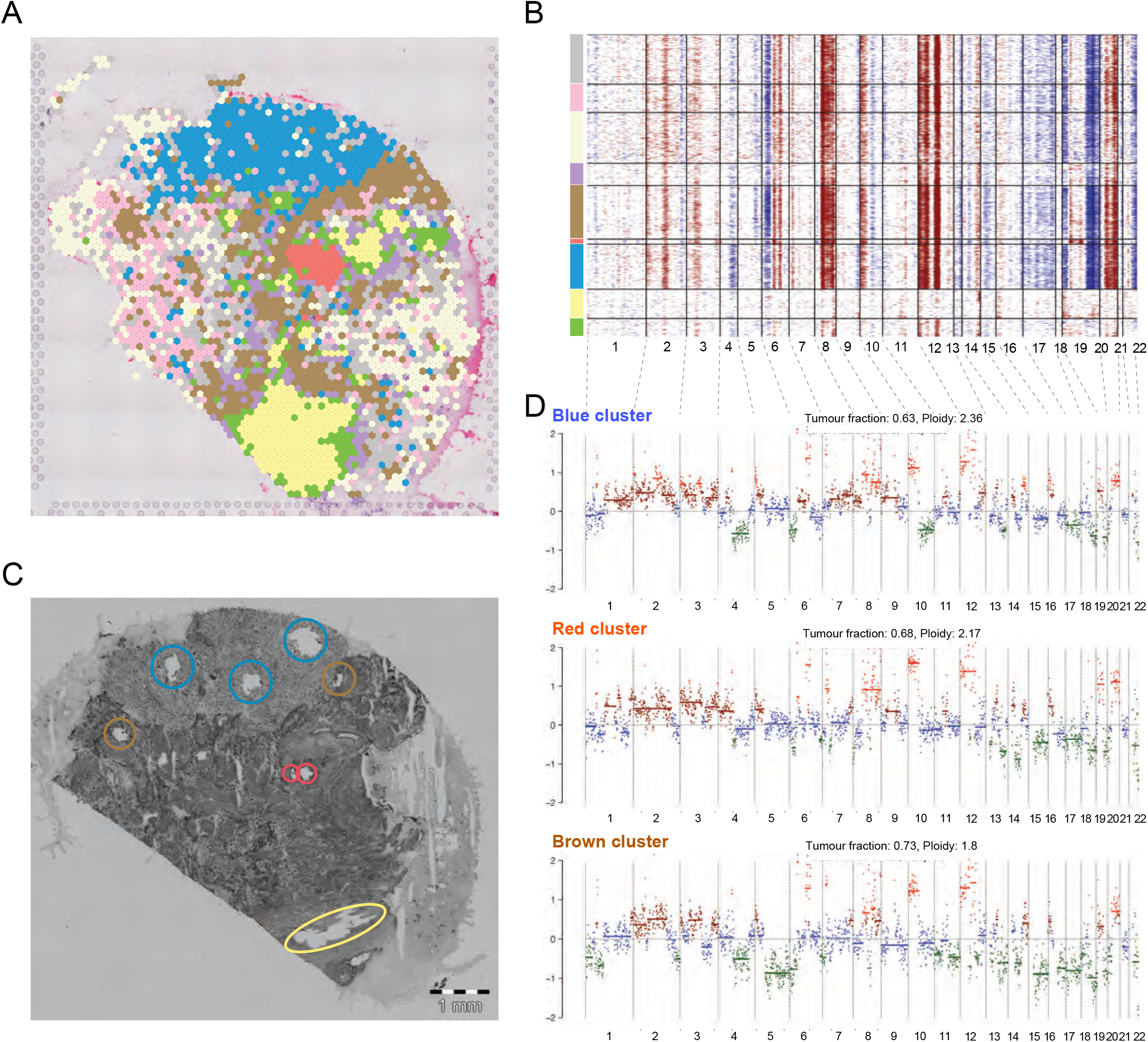
Copy number analysis reveals tumour subclones with spatially restricted patterns. **a**) Projection of spot clusters identified by inferCNV onto the tissue section for patient 1 **b**) Heatmap generated by inferCNV showing inferred CNA profiles of Visium spots for patient1. Horizontal black lines separate clusters identified by inferCNV; colours on the left correspond to colours used in panel (**a**). Red corresponds to predicted amplification, blue to predicted deletion. **c**) Adjacent tissue section adjacent from patient 1 showing areas collected for low pass whole genome sequencing. Colours of circles correspond to the colours of inferCNV clusters. **d**) ichorCNV CNA profiles of the blue, red, and brown malignant clusters. Green indicates 1 copy, blue 2 copies, brown 3 copies, red 4+ copies. Note, for the WGS of the blue, red, brown and non-malignant yellow clusters, 3, 2, 2 and 3 fragments were microdissected from each cluster, respectively. Tumour fraction and ploidy estimates from ichorCNV are indicated above each clone.

In addition to the multiple CNAs that were common across all predicted malignant inferCNV clusters in patient 1, we also observed that three of these clusters had large cluster-specific predicted CNAs. One cluster, localised to the top of the section (shown in blue in **Figure 2a**), has stronger evidence of deletion at chr4 than the other malignant clusters (**Figure 2b**). Similarly, the cluster shown in red (**Figure 2a-b**) has stronger evidence of amplification at chr19 and the corresponding spots are confined to an area in the middle of the tissue section. Lastly, the cluster shown in brown has stronger evidence of deletion at chr5 than the other malignant clusters (**Figure 2a-b**).

To validate the cluster-specific CNAs predicted for patient 1, we performed whole genome amplification and ultra-low-pass whole genome sequencing of microdissected regions corresponding to the putative blue, red, and brown tumour subclones and a non-malignant control region (yellow) identified in **Figure 2a-b** (**Methods**). For each cluster, multiple small tissue fragments were isolated from an adjacent section to that profiled by Visium (**Figure 2c**). We used ichorCNV ^36^ to pinpoint large-scale copy number alterations in the DNA isolated from each region. This approach validated both the CNAs predicted to be common to all tumour spots (large amplifications at chromosomes 8, 12, and 20, and the deletions at chromosomes 6 and 19, **Figure 2d**) and the subclone-specific alterations (the chr4 deletion, chr19 amplification, and chr5 deletion in the blue, red, and brown malignant clusters, respectively, **Figure 2d**). Thus, we concluded that the blue, red and brown areas of the tissue section shown in **Figure 2a** contain tumour subclones which are closely related with many shared CNAs but have acquired additional specific CNAs. A supplementary analysis comparing ichorCNV signals from different fragments in the same cluster highlighted likely further clonal heterogeneity in the blue, red, and brown clusters (**Supplementary Figure S6**).

For all eight samples, inferCNV predicted CNAs in regions that co-located with the Giotto tumour signal (**Figure1 c-d**). For six samples, inferCNV analysis predicted multiple potential tumour subclones (**Supplementary Figures S7, S9-S12**). For the remaining two sections, there were only small malignant areas with homogeneous CNAs predicted (**Supplementary Figures S8, S13**). Notably, there were few common alterations between patients (**Supplementary Figure S14**). For instance, the chr8:42541155-143878464, chr12:66765-47833132 and chr12:55817919-71667725 amplifications were only observed in patient 1 while the chr1:923928-15220480, chr1:23743155-28769775 and chr3:138944224-197956610 amplifications were specific to patient 3.

We next compared regions predicted to be amplified by inferCNV to recurrent amplifications identified in the 579 HGSOC tumour genomes sequenced by the TCGA (data was accessed from cBioPortal ^37^). Significantly, 188 of the 469 genes recurrently amplified in at least 20% of the TCGA HGSOC genomes overlap the 1,742 genes predicted to be amplified in at least one of our Visium samples (hypergeometric test p-value of 1.4*10^-35^). Similarly, 7 of 10 cytobands recurrently amplified in at least 20% of the TCGA HGSOC genomes were also predicted as amplified in at least one of our samples (**Supplementary Table ST3**). Our results extend previous studies on the polyclonality of HGSOC, revealing that even within a small area of <6.5mm^2^, multiple tumour subclones can be observed. Capturing this level of genetic heterogeneity will be a critical challenge for designing disease relevant cell, organoid and engraftment models of HGSOC.

### Different subclones from the same tumour can match different molecular subtypes

In 2008, Tothill *et al*.^24^ used bulk gene expression to describe four molecular HGSOC subtypes observed in pre-treatment samples: mesenchymal (C1.MES), immunoreactive (C2.IMM), differentiated (C4.DIF), and proliferative (C5.PRO) subtypes. Patients with C2.IMM and C4.DIF have been shown to have generally more favourable outcomes, while C1.MES and C5.PRO have poorer outcomes ^24,38^. However, there is still no consensus on their reproducibility and clinical significance ^39,40^. Most recently, a 55 gene predictor, the PrOTYPE (predictor of high-grade serous ovarian carcinoma molecular subtype) assay ^38^, has been developed to classify treatment-naïve primary tubo-ovarian HGSOC samples into these molecular subtypes. We used Seurat ^27^ to calculate module scores (*AddModuleScore* function) for each of these subtypes across the Visium slides using lists of subtype-enriched genes identified by PrOTYPE. This revealed that several subtypes were predicted to co-exist within several of our Visium sections (**Figure 3, Supplementary Figures S7-13, S15**) and that different inferCNV clusters within the same sample may correspond to different subtypes associated with different clinical outcomes. For example, the red and the blue inferCNV clusters observed in patient 5 (**Figure 3a**) have scores indicative of good outcome (high C4.DIF and low C1.MES), whereas the brown, grey, pink and purple clusters, in contrast, have low C4.DIF and high C1.MES scores (**Figure 3b,c**). Similarly, the brown and red clusters observed in patient 2 have lower C1.MES and higher C4.DIF scores than other clusters (**Supplementary Figure S7**). Notably, non-malignant areas consistently had higher C1.MES and lower C4.DIF scores. Our results suggest that classification of a tumour based on a bulk expression profile into the four previously described subtypes will depend highly on the area of tumour taken, the combination of subclones sampled and the patterns of infiltration by non-malignant cell types.

**Figure 3.**
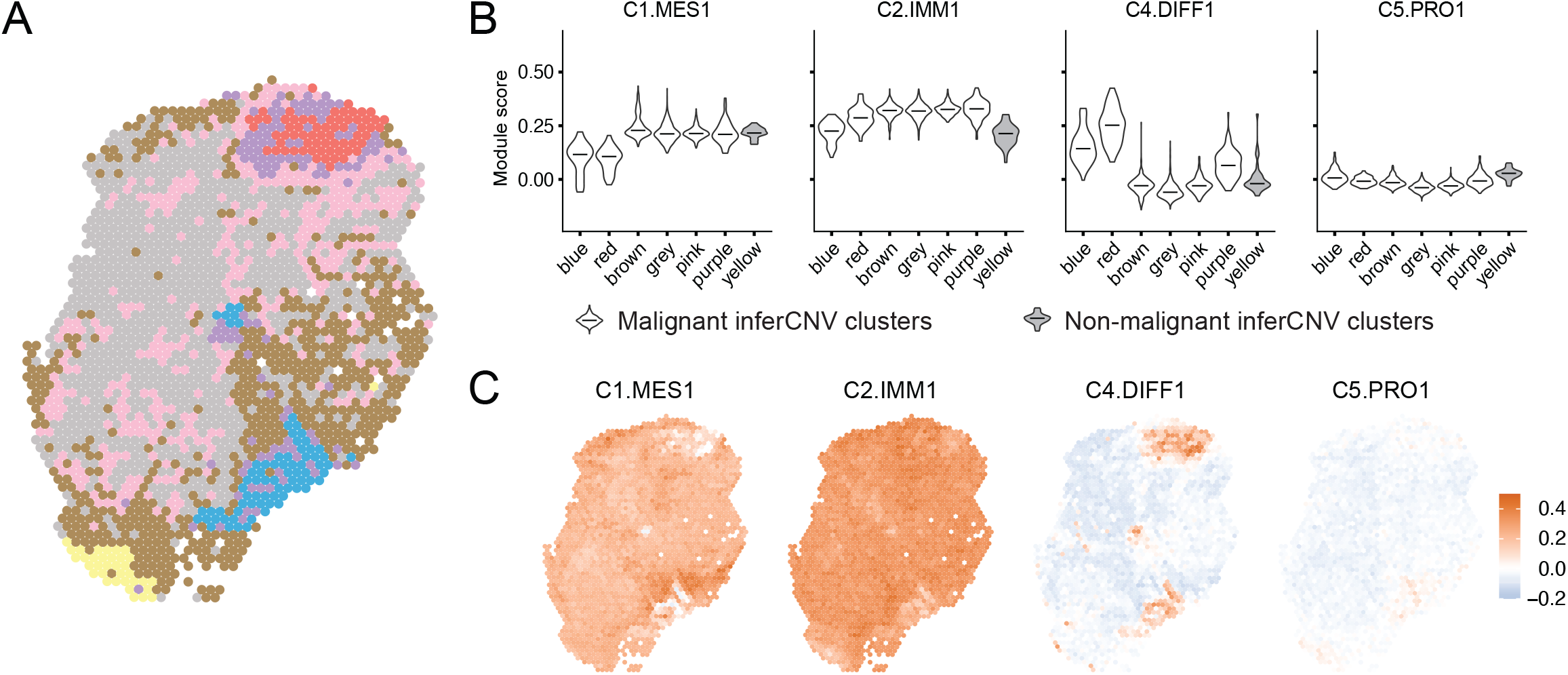
Intra-tumour heterogeneity observed in patient 5. **a**) Map of inferCNV clusters; yellow corresponds to normal tissue, other colours correspond to different malignant inferCNV clusters. **b, c**) Module scores calculated using *AddModuleScore* function in Seurat for genes overexpressed in four HGSOC molecular subtypes; shown in inferCNV clusters with median values indicated (**b**) and for each spot on the Visium slide (**c**).

### Gene expression differences between tumour subclones

InferCNV analysis predicted multiple malignant clusters for six of the eight samples profiled. To evaluate their heterogeneity, we first performed differential gene expression analysis between all pairs of predicted malignant clusters identified within each sample and merged together clusters with fewer than ten differentially expressed genes between them. In patient 7, this resulted in all four malignant inferCNV clusters being merged together. For the remaining five samples, between two and five clusters containing malignant subclones were obtained after merging. We then repeated the differential gene expression analysis for these merged clusters and the resulting tables shown in **Supplementary Table ST4** were used for the analyses described below.

Across the five samples, between 257 and 873 genes differentially expressed between subclone-containing clusters were identified (**Supplementary Table ST5)**. Notably, four of these gene lists were enriched for genes encoding cell surface or secreted proteins ^41^ (hypergeometric test p-value < 0.05), which suggests that these tumour subclones have different tumour microenvironments. As expected, comparing the differentially expressed genes with cell type specific expression from our scRNA-seq dataset showed that both altered gene expression between cancer cell subclones and differences in infiltration with non-malignant cell types contribute to differential expression between malignant inferCNV clusters (**Supplementary Tables ST4-5**). Examining the expression of the combined list of all 1,495 subclone differentially expressed genes in the scRNA-seq data revealed 41% (609 genes) were most highly expressed in tumour cells and 58% were most highly expressed in other cell types (55 genes in T cells, 44 in B/Plasma cells, 209 in macrophages, 94 in endothelial cells, 105 in mesothelial cells, 63 in myofibroblasts, 32 in fibro1, 23 in fibro2, 66 in fibro3, 34 in fibro4, and 144 in fibro5). The remaining 1% (17 genes) were not detected in our scRNA-seq dataset. Notably, many of the tumour cell differentially expressed genes were within subclone specific CNAs (**Supplementary Figure S16-20**). For example, eight of the tumour expressed genes (*FBN3, NDUFA7, ILF3, PRDX2, NR2F6, SUGP2, LPAR2, PDCD5*) from the red subclone of patient 1 are located in the validated red subclone-specific amplification on chr19 (**Supplementary Figure S16**).

We next examined the subclones of patient 1 in more detail. A total of 257 genes were differentially expressed between any of the red, purple, brown, blue or the merged grey_beige_pink clusters and 71 of these were most highly expressed in tumour cells (**Supplementary Table ST4**). We identified 83 genes most highly expressed in the predicted blue subclone (defined as being differentially up-regulated in the blue subclone in at least one of the pairwise comparisons and having the highest expression level in the blue subclone when compared to all other subclones). Of these, 23 are most highly expressed in tumour cells and 58 have the highest expression in other (non-malignant) cell types, indicating the blue subclone tumour microenvironment has higher levels of macrophages (18 genes) and myofibroblasts (9 genes). In contrast, 43 of the 91 genes more highly expressed in the red cluster were most highly expressed in tumour cells and only two and three genes were from macrophages or myofibroblasts, respectively. Notably, amongst the genes more highly expressed in the blue cluster, both tumour expressed genes (*CHI3L1, CLU, SLPI*) and genes from other cell types (*GPNMB, MGP, CRYAB, GPX3, MFAP4*) have been previously associated with poor prognosis and chemotherapy resistance in ovarian or other cancer types ^42–53^.

Taken together, our results show that the gene expression differences between different tumour subclone-containing clusters are explained by both the differences in tumour cell intrinsic expression as well as different patterns of immune and stromal cell infiltration. For the tumour cell intrinsic genes expressed at different levels in different subclones we find evidence that at least a subset is located directly within regions of copy number alteration. Finally, we find that the differentially expressed gene lists are enriched for cell surface or secreted proteins, which suggests that tumour subclones can have different tumour microenvironments.

### Cell-to-cell communication underlying different infiltration patterns

Given the enrichment of genes encoding secreted and plasma membrane proteins in the previous analysis, we next sought to examine whether differences in ligand expression in tumour cells and their interactions with cognate receptors expressed on non-malignant cell types might explain the different subclone-specific infiltration patterns predicted by Giotto. For this analysis, ligands and cognate receptors from connectomeDB2020 were used ^54^.

For the five samples with subclones, we first extracted all ligands that were differentially expressed between the subclones. This yielded 45, 38, 14, 109 and 84 subclonally differentially expressed ligands from patients 1, 2, 4, 5 and 6, respectively. We next annotated each ligand as most likely tumour-derived or microenvironment-derived using our scRNA-seq data based on a cell type with the highest expression level. After removing those with the highest expression in a non-malignant cell type, 9, 11, 4, 25, and 24 putative tumour intrinsic ligands remained, respectively, that were most highly expressed in tumour cells or plausibly subclone-specific (as they were not detected in any of the cell types identified in the scRNA-seq dataset) (**Supplementary Table ST6**). Importantly, expression of these ligands in a larger recently published metastatic ovarian cancer scRNA-seq dataset ^25^ further confirms their tumour cell specificity (**Supplementary Figure S21**).

To determine whether some of these ligands may influence the different non-malignant cell infiltration patterns reported by Giotto (**Supplementary Figures S5, S22**), we first calculated correlation between expression levels of each ligand and Giotto enrichment scores for each cell type across all malignant spots in each of the five samples (**Supplementary Table ST7**). The expectation for this analysis was that tumour cell derived ligands would be i) correlated with the Giotto tumour cell enrichment scores and ii) correlated or anticorrelated with the Giotto enrichment scores of non-malignant cell types for which the ligand directly or indirectly influenced their infiltration patterns. Reassuring us of the approach, expression of all but two ligands (*FGF19* in patient 1 and *ANXA1* in patient 5) was positively and significantly (FDR < 0.05) correlated with the Giotto tumour cell enrichment scores in at least one sample. This analysis also revealed many non-malignant cell infiltration patterns that were correlated or anticorrelated with tumour cell derived ligand expression levels. For example, B/Plasma cell infiltration was strongly anticorrelated with *CD24, FGF19, CD9* and *LAMA5* expression, T cell infiltration was correlated with the expression of *SLPI*, while macrophage infiltration showed anticorrelation with *FGF19* and positive correlation with multiple ligands (*SLPI, SLURP1, CD9, LCN2, L1CAM, CD24*) in patient 1.

To identify ligand-receptor pairs that may directly regulate the different infiltration patterns observed in **Supplementary Figure S22,** we selected ligands with significant correlation to a Giotto cell type enrichment score in **Supplementary Table ST7** and examined whether any cognate receptor was expressed in the same cell type. For the rest of the manuscript, we only focus on ligand-receptor pairs where the correlation between ligand expression and Giotto enrichment score of a target cell type is ≥ 0.1 or ≤ −0.1 (and significant with FDR < 0.05) and a cognate receptor for the ligand is detected in at least 10%of the cells of the same cell type (**Supplementary Table ST8**).

In total, 17 ligands were significantly correlated or anticorrelated with at least one non-malignant cell type expressing a cognate receptor and 37 tumour ligand-target cell pairs were observed across the five patients (**Table 1**). Of these tumour-derived-ligand-target-cell pairs, 26 were only observed as significant in one patient, four were significant and consistently correlated or anticorrelated in different patients, and seven (involving *L1CAM, LAMA5, SLPI* and *MMP7*) were correlated in some patients but anticorrelated in others.

**Table 1.** Correlations between tumour derived ligands and infiltrating cell types expressing cognate receptors. Summarises putative tumour intrinsic ligands (that were most highly expressed in tumour cells or plausibly subclone specific); non-malignant cell types with their respective Spearman correlation coefficients between Giotto enrichment score and ligand expression (filtered at ≥ 0.1 or ≤ −0.1 with FDR < 0.05); receptors detected in at least 10% of the target cells. Patient numbers are indicated in brackets.

| Ligand | Target cell | Receptors in target | Correlations (patient) | Anticorrelations (patient) |
| --- | --- | --- | --- | --- |
| ANXA1 | Macrophages | FPR1 | 0.3 (P5) |  |
| ANXA2 | Endothelial cells | ROBO4 | 0.12 (P6) |  |
|  | Macrophages | TLR2 | 0.52 (P5), 0.15 (P6) |  |
| BMP7 | Endothelial cells | BMPR2, ENG |  | -0.12 (P6) |
|  | Fibro5 (FN1, COL3A1) | BMPR2, ENG |  | -0.24 (P6) |
| CD24 | Endothelial cells | SELP |  | -0.13 (P2), -0.12 (P1) |
| CD47 | Macrophages | SIRPA |  | -0.16 (P2) |
|  | Mesothelial cells | SIRPA | 0.1 (P5) |  |
| CDH1 | B/Plasma cells | ITGB7 |  | -0.13 (P6) |
| CP | Mesothelial cells | SLC40A1 | 0.13 (P2) |  |
| CXCL10 | B/Plasma cells | CXCR3 | 0.16 (P5) |  |
|  | T cells | CXCR3 | 0.26 (P5) |  |
| FGF19 | Fibro5 (FN1, COL3A1) | FGFR1 |  | -0.1 (P1) |
|  | Mesothelial cells | FGFR1 | 0.12 (P1) |  |
| HMGB1 | Macrophages | HAVCR2, THBD |  | -0.18 (P5) |
|  | Myofibroblasts | THBD | 0.14 (P5) |  |
| L1CAM | Endothelial cells | CD9, ITGA5 | 0.1 (P1) |  |
|  | Fibro5 (FN1, COL3A1) | CD9, CNTN1, ITGAV |  | -0.2 (P6) |
|  | Macrophages | CD9, ITGA5 | 0.25 (P1) | -0.24 (P5) |
|  | Myofibroblasts | CD9 | 0.15 (P1), 0.19 (P5) | -0.11 (P6) |
|  | T cells | CD9 | 0.12 (P5) | -0.11 (P6) |
| LAMA5 | Endothelial cells | BCAM |  | -0.12 (P2) |
|  | Myofibroblasts | BCAM | 0.23 (P5) | -0.12 (P2) |
| LTBP3 | Fibro5 (FN1, COL3A1) | ITGB5 | 0.21 (P6) |  |
| MMP7 | B/Plasma cells | CD44 |  | -0.18 (P5), -0.14 (P4) |
|  | Fibro5 (FN1, COL3A1) | CD151, CD44 |  | -0.13 (P5) |
|  | Macrophages | CD151, CD44 | 0.14 (P4) |  |
|  | Mesothelial cells | CD151 | 0.11 (P5) | -0.14 (P4) |
|  | T cells | CD44 |  | -0.12 (P5) |
| RELN | Macrophages | ITGB1 |  | -0.14 (P5) |
|  | T cells | ITGB1 | 0.13 (P5) |  |
| SEMA3B | Fibro5 (FN1, COL3A1) | NRP2 |  | -0.1 (P6) |
| SLPI | Endothelial cells | PLSCR1, PLSCR4 | 0.13 (P1) | -0.16 (P2) |
|  | Fibro5 (FN1, COL3A1) | PLSCR1, PLSCR4 |  | -0.23 (P2) |
|  | Macrophages | PLSCR1 | 0.35 (P1) | -0.44 (P5), -0.24 (P2), -0.12 (P6) |
|  | Mesothelial cells | PLSCR1, PLSCR4 | 0.13 (P2) |  |
|  | Myofibroblasts | PLSCR1 | 0.2 (P1), 0.23 (P5) |  |

Several of these tumour ligand-target cell predictions are already implicated in HGSOC prognosis and tumour infiltration patterns. For example, in patient 5, *CXCL10* ligand expression strongly correlates with T cell scores and more weakly with B/Plasma cell scores and both cell types express the cognate receptor *CXCR3* (**Figure 4a-c, f**). We further examined the expression of *CXCL10* and *CXCR3* in a recently published scRNA-seq dataset comprising 51,786 cells from metastatic ovarian cancer samples ^25^, where, similarly to our dataset, *CXCL10* was detected in multiple cell types, but appeared more prevalent in the tumour cells, while *CXCR3* was detected mostly in T, NK, and dendritic cells (**Supplementary Figure S23**). Notably, *CXCL10* is a chemoattractant and its expression correlates with tumour infiltrating lymphocytes (TILs) in HGSOC and doubled overall survival ^55^. Similarly, in patient 1 and patient 2, *CD24* expression is anticorrelated with the Giotto scores for endothelial cells which express the cognate receptor *SELP*. In support of this anticorrelation, knockout of *CD24* in mice results in increased neovascularization of retina revealing a possible inhibitory role of *CD24* in angiogenesis ^56^. Notably, high cytoplasmic *CD24* expression is associated with poor prognosis of HGSOC ^57^.

**Figure 4.**
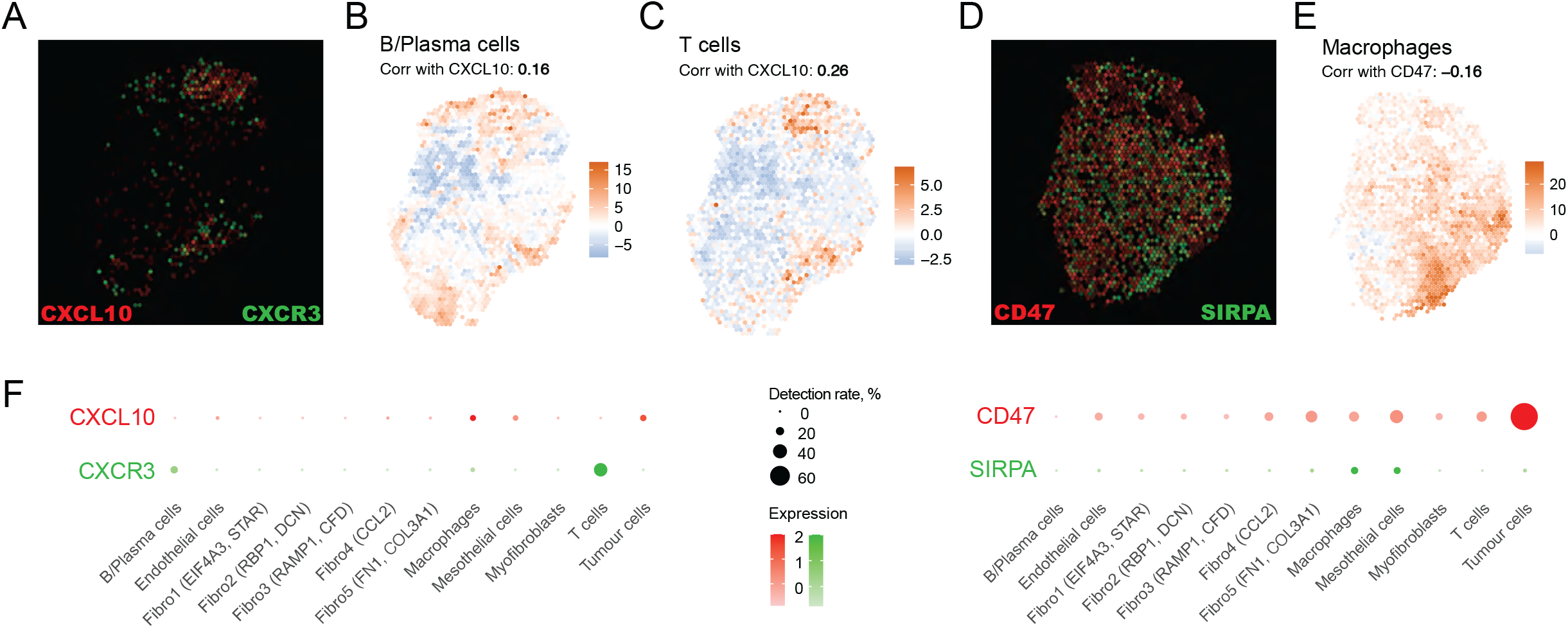
Selected ligand-receptor pairs potentially involved in maintaining tumour microenvironment. **a**) Overlayed expression of CXCL10 (red) and CXCR3 (green) in Visium data from patient 5. **b, c**) Giotto cell type enrichment scores for B/Plasma and T cells in Visium data from patient 55. **d**) Overlayed expression of CD47 (red) and SIRPA (green) in Visium data from patient 2. **e**) Giotto cell type enrichment scores for macrophages in Visium data from patient 2. **f**) Expression of these ligands and receptors in the scRNA-seq dataset.

For patient 2, *CD47* expression is significantly anticorrelated with the Giotto scores for macrophages which express the cognate receptor *SIRPA* (**Figure 4d-f**). Similarly, in the Zhang *et al*. dataset ^25^, *CD47* was more specific to tumour cells (**Supplementary Figure S23**). *CD47* over-expression in HGSOC has been previously reported as potentially associated with poor prognosis ^58,59^. *CD47* plays an important role by sending an antiphagocytic signal via *SIRPA* to tumour associated macrophages which can be targeted by anti-CD47 agents ^60^. Consistent with the anticorrelation observed in our analysis, anti-CD47 monoclonal antibody treatment was shown to increase macrophage infiltration of tumours in a HGSOC xenograft model ^61^.

Taken together, our results indicate that tumour subclones are prevalent in HGSOC and that they may directly influence the infiltration patterns of non-malignant cell types by differential expression of ligands.

## Discussion

Here, we have used Visium spatial transcriptomics to study tumour heterogeneity in HGSOC samples from eight patients who underwent platinum and taxane-based neoadjuvant chemotherapy. Copy number inference helped delineate tumour regions from non-malignant areas but also revealed subclonal heterogeneity in five of the eight samples studied. Our work mirrors a recent report of spatially restricted subclones observed via spatial inferred CNVs (siCNVs)^62^. For one sample, microdissection and ultra-low-pass whole genome sequencing confirmed the presence of subclone-specific CNAs. Finding evidence of multiple subclones in small 6.5mm^2^ sections suggests there is an even higher degree of intratumoural subclonal heterogeneity than previously appreciated. Seeking to independently validate this observation we performed inferCNV analysis of Visium data from a complementary study on 12 treatment naive HGSOC samples ^63^. Patient-specific CNAs were clearly identified but no subclones were predicted (**Supplementary Figure S24**). The most likely explanation for the discrepancy is that the sections in the other study were too small to sample multiple tumour clones (a median of 364 spots per sample) in comparison to our sections (a median of 2,507 spots per sample). Notably, Schwarz *et al*. have reported intratumoural clonal heterogeneity is present in pre-treatment samples and correlates with post-chemotherapy survival outcomes of HGSOC patients ^8^. Using pre- and post-treatment samples, they find evidence of clonal expansion and that patients with higher clonal expansion had poorer outcomes. Zhang *et al*. similarly observed evidence of clonal expansion in neoadjuvant chemotherapy treated samples with selection for tumour cells with what they refer to as a high stress-associated score which they show is associated with poor progression-free survival using deconvoluted RNA-seq data of a larger TCGA cohort ^25^. Notably, many of the subclonally differentially expressed genes that we identified have previously been associated with HGSOC prognosis and chemotherapy response. This reinforces the position that subclones are prevalent in HGSOC, that they have differential sensitivity to chemotherapy and, thus, may lead to recurrence and relapse.

We also find that, in addition to genes differentially expressed by tumour subclones, approximately 58% of genes differentially expressed between the inferCNV clusters containing these subclones are from non-malignant cell types (the most abundant being 20% from fibroblasts and 14% from macrophages). This is also reflected in the substantial differences in the infiltration patterns observed from the Giotto analysis (**Supplementary Figure S22**). These analyses show that different tumour subclones can have different microenvironments which can correspond to different previously reported molecular subtypes (**Figure 3**). This builds upon observations by Schwede *et al*. and Izar *et al*. that these subtype signatures are more a reflection of stromal content rather than underlying tumour cell differences ^40^ with the immunoreactive and mesenchymal subtypes indicative of the abundance of immune infiltrates and fibroblasts respectively rather than distinct subsets of malignant cells ^19^.

Next our focus turned to why different sets of non-malignant cell types are associated with subclones that have different copy number alterations. Our analyses revealed an over-representation of plasma membrane and secreted proteins in the sets of subclonally differentially expressed genes. Examining this further, we identified sets of ligands differentially expressed by subclones that were correlated or anticorrelated with infiltration of a non-malignant cell type. We then highlighted a subset of ligands where the cognate receptor was expressed in the same non-malignant cell type suggesting the tumour subclone may directly influence the observed infiltration pattern (**Table 1, Figure 4**). Others have previously attempted to relate tumour-expressed ligands to differences in infiltration patterns ^64–66^; however, to our knowledge, this is the first study to demonstrate that subclones express different ligands which then influence non-malignant cell infiltration. Notably, these associations were largely patient- or subclone-specific. For example, the correlation between tumour-derived *CXCL10* and infiltration of *CXCR3*-expressing T cells and B/plasma cells was only observed in subclones of one patient (**Figure 4**). A similar association between *CXCL16* expressing tumour cells and *CXCR6* expressing T cells has been reported in patients with highly infiltrated tumours ^22^.

In closing, there are several limitations of our study that will need to be addressed in the future using new technologies and by studying multiple tumour regions from larger cohorts of patients. The first is that the current resolution of the Visium spatial technology is not at the level of a single cell, thus, we needed to infer which cell types are present under a spot based on projection of marker genes from scRNA-seq data. With single-cell resolution cDNA-based spatial transcriptomics (e.g Slide-seq ^67^, STOmics ^68^, VisiumHD), it will become possible to directly identify which cells are expressing these genes. The second limitation is that we have inferred copy number alterations and subclones based on transcriptome data. Methods that directly provide spatial genomic information (e.g., Slide-DNA-seq ^69^) would improve our ability to call smaller alterations and single nucleotide variants missed in our data. The last limitation is the size of the cohort considered. There was little overlap between CNAs observed in each of the eight patients and only modest overlap with CNAs reported by the TCGA. For five of the patients, we confidently observed subclones with different CNAs. Non-malignant infiltration patterns of the tumour regions varied substantially between patients and subclones. For example, substantial infiltration by T cells and neutrophils was only observed in subclones of one patient while subclones with strong or weak macrophage infiltration were observed for all patients with subclones. Similarly, we observed subclone-specific ligand expression patterns (e.g., *CXCL10* in subclones of one patient).

Our results further highlight the high degree of interpatient and intrapatient heterogeneity seen in HGSOC. This heterogeneity is clinically important as it likely explains resistance and recurrence even after an initial good response. It will be critical moving forward that we better capture the subclonal populations present in each patient and relate that back to their tumour microenvironments and ultimately drug sensitivities and prognosis.

## Methods

### Ovarian tumour samples and consent

High-grade serous ovarian tumours from eight patients diagnosed with stage III-IV cancers were included in this study. Patients were treated with 3-6 cycles of platinum-based chemotherapy. All tumour samples were derived from ovarian sites during interval debulking surgery. Fresh tumours were either collected in RPMI (ThermoFisher Scientific) supplemented with penicillin and streptomycin (Sigma) for single-cell dissociation or immediately snap-frozen and stored in −80°C tumour bank until retrieved for Visium experiments.

The chemotherapy response scores (CRS, 3 tier) for each patient were determined as previously described (PMID: 26124480) by assessing the largest (macroscopic) omental tumour deposit for features of regression based on the following: score 1 (no/minimal tumour response), score 2 (partial tumour response) and score 3 (complete/near complete response, cell groups measuring <2mm each, or no residual tumour) ^26^.

The study was approved by St John of God Health Care (SJGHC), The University of Western Australia (UWA) and Curtin University Human Research Ethics Committees (#1217 and RA/4/20/5784). All participants were given information about the study and provided written informed consent before enrolment.

### Single-cell suspension for scRNA-seq

HGSOC tumour were dissociated using the tumour dissociation kit 2, human, from Miltenyi Biotec [130-095-929] as per manufacturers’ instruction. Tumour tissue (0.2 – 1 g) was cut into small pieces of 2-4 mm and placed into a gentleMACS C-tube [Miltenyi Biotech; 130-093-237] containing the enzyme mix from the kit. The tube was then placed onto the gentleMACS octo dissociator (Miltenyi Biotech) and processed using the 37C_h_TDK_1 program with the associated incubation times indicated in the protocol. Complete tissue dissociation was confirmed by the absence of visible tissue chunks. The resulting tumour homogenate was filtered using a 70-μm MACS SmartStrainer and washed with RPMI. Cell suspension was further filtered through 40-μm strainers to remove cell clumps. The viability was assessed by ReadyProbe Cell Viability Imaging Kit (ThermoFisher Scientific) to ensure the viability was >90%.

### Single-cell RNA-seq profiling

Cryostored cells were rapidly thawed in a water bath set at 37 C. 1mL of Media (RPMI1640 +10%FBS) was then added to the cells, which were then mixed and transferred to a 15mL falcon tube. The cryovial was then rinsed with another 1mL of media which was subsequently added to the 15mL falcon tube in a dropwise fashion. 7mL of media was then added to the falcon tube dropwise using a serological pipette. The cells were then centrifuged at 300g for 5 minutes. The supernatant was removed, leaving behind 1mL of media, and another 1mL of media was added and the cells were resuspended. 2mL of DPBS+0.04%BSA was then added to the cells, followed by another centrifugation at 300g for 5 minutes. The supernatant was then removed and the cells were resuspended in 1mL of DPBS+0.04%BSA and then subsequently run through a 40uM filter. Cells were then counted and viability was checked using the Countess II automated cell counter and the readyprobes red/blue viability kit (Thermo Fisher Scientific). Libraries were prepared in accordance with the protocol for 10x Chromium Single Cell 3’ v2 (10x Genomics). Sequencing was performed on a NovaSeq 6000 (Illumina).

### Visium spatial transcriptomic profiling

Frozen tissue fragments were embedded in Tissue-Tek O.C.T. Compound (25608-930, VWR) according to the Visium Spatial Protocols – Tissue Preparation Guide (CG000240 Rev A, 10x Genomics) and stored immediately at −80°C until further use. Hematoxylin and eosin staining of 10μm cryosections from each O.C.T. block were assessed by a pathologist to confirm tissue type and tumour content. Samples with adequate tumour content were selected for use in the gene expression workflow.

To assess the quality of the selected tissue blocks, RNA was isolated from serial sectioned tissues totalling 80μM thickness and its RNA integrity number (RIN) was calculated using the Agilent 4200 TapeStation system. Samples which had a RIN ≥7 were considered good quality and selected to proceed with the experiment. Each Visium Spatial Gene Expression Slide (2000233, 10x Genomics) was used to analyse up to four tissue samples, i.e. one section per sample block. Of the four samples, one block was randomly selected for tissue optimisation using a Visium Spatial Tissue Optimisation Slide (3000394, 10X Genomics). Serial tissue sections at 10μM thickness were placed on seven capture squares of the pre-chilled tissue optimisation slide with the remaining square left empty. Different tissue permeabilization times were tested on 6 of the sections at 10-minute intervals to a maximum of 60 minutes. The remaining tissue section represented a negative control for permeabilisation while the empty well served as a positive control to which a reference RNA was added (QS064, Life Technologies). The optimal permeabilisation time point was 30 minutes and was therefore used as the permeabilisation time on the gene expression samples.

A Nikon Eclipse Ni-U microscope with a 10x objective in large scale imaging mode (Nikon, NIS-Elements AR 5.21.00) was used to take brightfield images of the Visium Spatial Tissue Optimisation and Gene Expression slides. The same settings were used to collect fluorescent images of the optimisation slides via a Texas Red HYQ filter cube at 1.5 seconds exposure time. Images were automatically stitched via blending with a 10% tile overlap. Original files were saved as the default ‘.ND2’ format and exported to ‘.tiff’ or ‘.jpeg’ using NIS-Elements AR or ImageJ, respectively.

Libraries were prepared according to the Visium Gene Expression User Guide (CG000239, Rev A, 10X Genomics) and pooled to a final library concentration of 1.8nM. The samples were loaded on a NovaSeq 6000 System (Illumina) using NovaSeq 6000 SP Reagent kit (200 cycles, 20040326, Illumina) and sequenced at a depth of approximately 150M reads per sample. The read protocol was set as the following: read 1 at 28 cycles, i7 index read at 10 cycles, i5 index read at 10 cycles and read 2 at 120 cycles.

Manual image alignment and spot selection of the H&E brightfield images was performed in the Loupe Browser.

### Ultra-low-pass DNA sequencing

Frozen tissue was sectioned (10 μm) and mounted to standard superfrost slides, methanol fixed, stained with hematoxylin and eosin, and scanned on a CellCelector (ALS). Using the CellCelector, the long edge of a 150 μm glass capillary was used to mechanically scrape small tissue sections from the slide which were aspirated and deposited in 1 μL of PBS (10 mM Phosphate, 2.68 mM Potassium Chloride, 140 mM Sodium Chloride, 18912014, Thermo Fisher Scientific) in 0.2 mL PCR tubes (Eppendorf). Tissue sections were subjected to whole genome amplification using the Ampli1 WGA Kit (Silicon Biosystems) to the manufacturer’s instructions. Following amplification, 400 bp sequencing libraries were constructed using the Ampli1 Low-Pass Whole Genome Sequencing Kit for Ion Torrent (Silicon Biosystems) to the manufacturer’s instructions. Libraries were diluted to 50 pM, loaded into an Ion Chef for template preparation and loading into an Ion 530 chip, and then sequenced for 525 flows on an Ion S5 (Thermo Fisher Scientific). Sequencing data was aligned to hg38 and indexed using Torrent Server (V 5.16) with depths ranging from 0.1 to 0.3x. Following alignment and indexing, .wig files were generated using readCounter in 1 Mb windows from HMM Copy Utils [https://github.com/shahcompbio/hmmcopy_utils]. IchorCNA (v0.2.0)^36^ was used to detect somatic copy number alterations with 1 Mb bins and the run parameters set to “--ploidy “c(2,3,4)”, --normal “c(0.05)”, --includeHOMD False, --chrTrain “c(1:22)”, and --estimateScPrevalence False. The non-malignant yellow regions observed for patient 1 were used to construct a panel of normals using IchorCNA’s createPanelOfNormals.R script.

### Single-cell RNA-seq data analysis

#### scRNA-seq data processing

FASTQ files were processed using Cell Ranger 3.0.2 with refdata-cellranger-GRCh38-3.0.0 reference. Raw gene-barcode matrices from Cell Ranger output were used for downstream processing. Cells were distinguished from background noise using EmptyDrops ^70^. Genes detected in a minimum of 3 cells were retained; cells with at least 500 genes, at least 1000 UMIs and under 15% of mitochondrial reads were retained. Seurat v3 was used for sample normalisation (SCTransform, mitochondrial and ribosomal mapping percentage were regressed out), integration (anchor-based method with 3000 variable genes), dimensionality reduction and clustering (using first 30 principal components), and differential expression analysis (Wilcoxon test)^71^.

#### Inferring cell identity

To infer cell identities, we first performed a reference-based annotation using scMatch ^28^ and a reference dataset from Olbrecht *et al*. ovarian cancer samples ^21^. To construct the reference dataset, we obtained gene counts and cell types reported in ^21^; counts were normalised to cell library size and averaged within each cell type to derive reference vectors for scMatch. We then used scMatch with parameters *--testMethod s --keepZeros n* to label each individual cell with the closest cell type identity from the reference dataset. This resulted in seven major cell types: tumour cells, fibroblasts, ovarian stroma, endothelial cells, monocytes, T cells, and B cells. We then examined expression of differential expressed and cell type marker genes in these seven cell types and based on this relabelled two of them to better reflect the cell identity (B cells to B/Plasma cells based on the expression of *IGHG1, IGHG3, JCHAIN;* monocytes to macrophages based on the expression of *CCL3, CXCL8, HLA-DRA*) (**Supplementary Figure S2a, b**).

Cells labelled as ovarian stroma and fibroblasts formed multiple visually distinct cell groups. To explore possible subtypes, we extracted these cells from the dataset and reran principal component and clustering analysis for this subset. We identified ten clusters (**Supplementary Figure S2c, d**). Cell identities were assigned to these clusters based on their specific differentially expressed genes and cell type gene markers.

**Cluster 1** was characterised by high expression of contractile genes including *TAGLN, ACTA2, MYL9, MYH11, PLN* and was labelled Myofibroblasts. **Cluster 6** was labelled Mesothelial cells based on *CALB2, MSLN, SLPI, KRT8, KRT18* expression, as per Qian *et al*.^72^ and Olbrecht *et al*.^21^. **Cluster 3** showed high expression of *COL1A1, COL1A2, COL3A1, SPARC, FN1* and was labelled Fibro5 (*FN1, COL3A1*). In **Cluster 8,** *CFD* and *RAMP1* were the top DEGs, and the cluster was labelled Fibro3 (*RAMP1, CFD*) - these cells might correspond to adipogenic fibroblasts, as per Qian et al.^72^ and Olbrecht et al. ^21^. **Cluster 7** specifically overexpressed *CCL2* and we labelled it Fibro4 (*CCL2*). **Cluster 2** did not have genes strongly overexpressed with logFC > 1, but overexpressed *LUM, DCN, GSN* with logFC >= 0.5 and was labelled Fibro2 (*RBP1, DCN*). **Cluster 0** and **Cluster 5** were labelled Fibro1 (*EIF4A3, STAR*).

**Cluster 4** showed high expression of stress response-related genes, such as *HSPA6, HSPA1B, DNAJB1, HSPA1A*, hence we assumed it corresponded to cells showing strong stress response and removed it from the downstream analysis. Finally, **Cluster 9** had high expression of genes normally expressed in immune cells, such as *B2M, CCL5, HLA-A, HLA-B, CXCR4*, and these cells co-clustered with T cells in the superset, hence, we concluded this cluster corresponded to doublets and removed it from the downstream analysis.

#### Visium data processing

FASTQ files were processed using Space Ranger 1.0.0 with GRCh38-3.0.0 reference in the manual alignment mode. Filtered gene-barcode matrices from Space Ranger output were used for downstream analyses; barcodes with less than 400 genes were excluded. Seurat v3 was used for sample normalisation (SCTransform, mitochondrial and ribosomal mapping percentage were regressed out), individual sample clustering (using first 30 principal components), integration (anchor-based method with 3000 variable genes), dimensionality reduction and clustering (using first 30 principal components), and differential expression analysis (Wilcoxon test)^71^.

### CNA inference

#### Identification of background spots

Gene expression data from all eight Visium samples were analysed together using Seurat v3 ^71^. We performed normalisation using SCTransform ^73^ and regressed out the percentage of mitochondrial and ribosomal counts. Then an anchor-based dataset integration was performed with 3000 features, followed by clustering and UMAP projection using 30 first principal components. We identified seven top-level clusters at resolution 0.2 and used *FindMarkers* function to compute differentially expressed genes.

Manual curation of differentially expressed genes identified two clusters (clusters 2 and 6) corresponding to stromal tissues (expressing *DCN, TAGLN, ACTA2, VWF* and other markers, see **Supplementary Figure S25**). We next performed one more round of analysis to refine those two clusters. First, the spots corresponding to the two clusters were extracted into a separate candidate stromal dataset and normalisation, integration, and downstream analysis were repeated as above. In this candidate stromal dataset, we identified seven clusters at resolution 0.6. One of these clusters showed higher expression of cancer genes (Clusters 3, **Supplementary Figure S25d,e**) and the corresponding spots were excluded from the stromal dataset.

Initially, this stromal dataset was used as a background for CNA inference; however, in the downstream analyses we found that the predicted malignant and non-malignant tissue areas were intermixed (with no clear zonation) in two out of the eight tissue sections; one more tissue section had large necrotic areas. To reduce possible noise due to the background spot set used for CNA inference, we excluded spots corresponding to these three samples (patient 2, 6 & 7) from the stromal dataset and used the remaining 2,772 spots from the remaining 5 samples as a background for inferCNV inference reported here.

#### inferCNV

inferCNV ^35^ was run for each sample independently using stromal spots defined above as a background with the following parameters: *cutoff=0.1, denoise=T, HMM=F*. Spots in each sample were clustered using the default parameters and the dendrogram was split into clusters with visually distinctive CNA profiles.

### Cell type enrichment analysis

We used Giotto ^29^ to estimate cell type enrichment across different spots in each of the Visium samples. To identify gene sets for the enrichment analysis, we started with our scRNA-seq dataset. First, differentially expressed genes were calculated pairwise for each possible pair of the 12 cell types. This was done using the *FindMarkers* function in Seurat with the minimum detection rate threshold of 0.5. For each cell type, we then selected genes that passed the thresholds of logFC >= 0.5 and FDR < 0.05 in at least 10 out of the 11 pair-wise tests (i.e. genes that were significantly differentially over-expressed in that cell type when compared to at least 10 of the other cell types). We next removed genes that were identified this way for more than one cell type. This approach failed to identify suitable gene sets for four of the cell types (Fibro4 (*CCL2*) (identified 1 gene only), Fibro3 (*RAMP1, CFD*) (identified 4 genes only), Fibro2 (*RBP1, DCN*) and Fibro1 (*EIF4A3, STAR*) - no genes identified). We then manually curated the gene lists for T cells, B cells, and macrophages; we removed non-specific genes based on their expression in other cell types in the FANTOM5 dataset ^74^ and added alternative markers that should be specific to those cell types. We also included additional manually curated gene lists for neutrophils, mast cells and adipocytes. See **Supplementary Table ST9** for the final table with genes used for the cell type enrichment analysis and **Supplementary Figure S26** for the expression of these genes in our scRNA-seq dataset. Genes obtained in this manner were then used as input for the PAGE algorithm in Giotto, which calculated enrichment scores for the corresponding cell types for each Visium spot.

### TCGA recurrent alterations comparison

TCGA CNA data were downloaded from cBioPortal (www.cbioportal.org) from Ovarian Serous Cystadenocarcinoma (TCGA, Firehose Legacy) dataset. CNA genes reported for 579 samples were filtered using the frequency of 20% as a threshold. For this comparison, genes predicted to be amplified in our Visium samples were selected with the threshold of >= 1.1 imposed on inferCNV signal averaged across spots in each of inferCNV-based clusters. Note, deletions were not tested as recurrent deletions are rare (there are no regions or genes reported as recurrently deleted in at least 20% of the TCGA HGSOC genomes).

## Data availability

All data analysed within this manuscript are publicly available from the Gene Expression Omnibus (GEO) repository with the primary accession code GSE211956.

## Authors contributions

The project was conceived by ARRF, LdK, PAC, YY1. Computational analyses and data interpretation were carried out by ED. AT and MB did the pathological annotations. GM, SS carried out the surgical collections. SB handled the ethics.YY1 prepared the single cell suspensions. AB and EG did the low pass WGS and ichorCNV analysis. MJ, KH, DO’M, YY2 generated scRNA/snRNA-seq libraries. LdK generated the Visium libraries. MJ did the sequencing. The manuscript was written by ED and ARRF with input from YY1, PAC and all authors.

## Acknowledgements

We would like to acknowledge the patients who participated in this study. We would like to acknowledge Joost Lesterhuis for useful comments on the manuscript.

## Supplementary Materials

### Supplementary Tables

**Supplementary Table ST1**: Sample information for Visium and scRNA-seq samples.

**Supplementary Table ST2**: Cell type labels assigned to individual cells in the scRNA-seq dataset.

**Supplementary Table ST3**: Cytobands recurrently amplified in at least 20% of the TCGA HGSOC genomes.

**Supplementary Table ST4**: Differentially expressed genes (DEGs) calculated between all possible pairs of putative malignant subclones identified within each Visium sample. Seurat *FindMarkers* function was used with logfc.threshold = 0.5, min.pct = 0.5 and otherwise default parameters.

**Supplementary Table ST5**: Summary for DEGs shown in Supplementary Table ST4.

**Supplementary Table ST6**: Ligands selected for each sample for cell-to-cell communication analysis.

**Supplementary Table ST7**: Correlation between expression levels of selected ligands and Giotto enrichment scores for each cell type, calculated across all malignant spots in each of five Visium samples.

**Supplementary Table ST8**: Selected ligand-receptor pairs in five Visium samples. Includes average expression of each ligand in each putative malignant subclone, correlation between ligand expression and Giotto enrichment scores for non-malignant cell types calculated across all malignant spots and filtered to have correlation coefficient of ≥ 0.1 or ≤ −0.1 with FDR < 0.05, average expression and detection rate of the cognate receptor in the same cell type.

**Supplementary Table ST9**: Gene signature table used for Giotto cell type enrichment analysis. Each column corresponds to one cell type, the value of 1 indicates that the corresponding gene was included for the corresponding cell type.

**Supplementary Table ST10**: Genes differentially expressed between malignant spots of Visium samples of CRS1 and CRS3 patients.

### Supplementary Figures

**Supplementary Fig. S1: Study design.** Samples were collected during interval debulking surgery from patients who underwent 3-4 cycles of platinum and taxane treatment. Eight samples were profiled using 10x Genomics Visium and five samples using 10x Genomics 3’ Gene Expression solution.

**Supplementary Fig. S2: Annotation of the single-cell RNA-seq HGSOC dataset. a)** UMAP showing top-level cell type annotations inferred by scMatch with Olbrecht *et al*.reference dataset. **b)** Differentially expressed genes (DEGs) identified in the top-level cell types; top DEGs based on the ratio of detection rates are shown for each cell type. **c)** UMAP showing subclustering of fibroblasts and stromal cells. **d)** DEGs identified in the subclusters shown in (**c**); top DEGs based on the ratio of detection rates are shown for each cell type.

**Supplementary Fig. S3: Fine-grain annotations of the scRNA-seq dataset. a)** UMAP showing 12 cell types. **b)** Top differentially expressed genes (DEGs) over-expressed in each of the cell types, ranked based on logFC. **c)** Top DEGs over-expressed in each of the cell types, ranked based on the ratio of detection rates. DEGs were calculated for each cell type vs all other cells.

**Supplementary Fig. S4: Correlation between Giotto cell type enrichment scores across all Visium samples and spots.**

**Supplementary Fig. S5: Giotto cell type enrichment scores for eleven cell populations in Visium samples.**

**Supplementary Fig. S6: Validation of inferCNV predictions using ultra-low-pass whole genome sequencing.** IchorCNV analysis was performed for each of the numbered regions separately, using tongue tissue as a background. IchorCNV CNA profiles are shown for each region, where green indicates 1 copy, blue 2 copies, brown 3 copies, red 4+ copies.

**Supplementary Fig. S7: Visium data summary for patient 2. a)** Heatmap generated by inferCNV showing inferred CNA profiles of Visium spots. Horizontal black lines separate clusters identified by inferCNV. Red corresponds to predicted amplification, blue to predicted deletion. **b)** Projection of spot clusters identified by inferCNV onto the tissue section. **c, d)** Module scores calculated using *AddModuleScore* function in Seurat for genes overexpressed in four HGSOC molecular subtypes; shown in malignant inferCNV clusters with median values indicated (**c**) and for each spot on the Visium slide (**d**).

**Supplementary Fig. S8: Visium data summary for patient 3. a)** Heatmap generated by inferCNV showing inferred CNA profiles of Visium spots. Horizontal black lines separate clusters identified by inferCNV. Red corresponds to predicted amplification, blue to predicted deletion. **b)** Projection of spot clusters identified by inferCNV onto the tissue section. **c, d)** Module scores calculated using *AddModuleScore* function in Seurat for genes overexpressed in four HGSOC molecular subtypes; shown in malignant inferCNV clusters with median values indicated (**c**) and for each spot on the Visium slide (**d**).

**Supplementary Fig. S9: Visium data summary for patient 4. a)** Heatmap generated by inferCNV showing inferred CNA profiles of Visium spots. Horizontal black lines separate clusters identified by inferCNV. Red corresponds to predicted amplification, blue to predicted deletion. **b)** Projection of spot clusters identified by inferCNV onto the tissue section. **c, d)** Module scores calculated using *AddModuleScore* function in Seurat for genes overexpressed in four HGSOC molecular subtypes; shown in malignant inferCNV clusters with median values indicated (**c**) and for each spot on the Visium slide (**d**).

**Supplementary Fig. S10: Visium data summary for patient 5. a)** Heatmap generated by inferCNV showing inferred CNA profiles of Visium spots. Horizontal black lines separate clusters identified by inferCNV. Red corresponds to predicted amplification, blue – predicted deletion. **b)** Projection of spot clusters identified by inferCNV onto the tissue section. **c, d)** Module scores calculated using *AddModuleScore* function in Seurat for genes overexpressed in four HGSOC molecular subtypes; shown in inferCNV clusters with median values indicated (**c**) and for each spot on the Visium slide (**d**). Note, panels **b**, **c**, **d** are duplicating Figure 3.

**Supplementary Fig. S11: Visium data summary for patient 6. a)** Heatmap generated by inferCNV showing inferred CNA profiles of Visium spots. Horizontal black lines separate clusters identified by inferCNV. Red corresponds to predicted amplification, blue to predicted deletion. **b)** Projection of spot clusters identified by inferCNV onto the tissue section. **c, d)** Module scores calculated using *AddModuleScore* function in Seurat for genes overexpressed in four HGSOC molecular subtypes; shown in malignant inferCNV clusters with median values indicated (**c**) and for each spot on the Visium slide (**d**).

**Supplementary Fig. S12: Visium data summary for patient 7. a)** Heatmap generated by inferCNV showing inferred CNA profiles of Visium spots. Horizontal black lines separate clusters identified by inferCNV. Red corresponds to predicted amplification, blue to predicted deletion. **b)** Projection of spot clusters identified by inferCNV onto the tissue section. **c, d)** Module scores calculated using *AddModuleScore* function in Seurat for genes overexpressed in four HGSOC molecular subtypes; shown in malignant inferCNV clusters with median values indicated (**c**) and for each spot on the Visium slide (**d**).

**Supplementary Fig. S13: Visium data summary for patient 8. a)** Heatmap generated by inferCNV showing inferred CNA profiles of Visium spots. Horizontal black lines separate clusters identified by inferCNV. Red corresponds to predicted amplification, blue to predicted deletion. **b)** Projection of spot clusters identified by inferCNV onto the tissue section. **c, d)** Module scores calculated using *AddModuleScore* function in Seurat for genes overexpressed in four HGSOC molecular subtypes; shown in malignant inferCNV clusters with median values indicated (**c**) and for each spot on the Visium slide (**d**).

**Supplementary Fig. S14: Summary of inferCNV signal predicted in eight Visium samples.** Each panel shows one chromosome. InferCNV signal was averaged for each sample across all malignant spots. Genomic areas with InferCNV signal above 1.1 were considered amplified for the between-sample and TCGA dataset comparisons.

**Supplementary Fig. S15: Visium data summary for patient 1.** Module scores calculated using *AddModuleScore* function in Seurat for genes overexpressed in four HGSOC molecular subtypes; shown in malignant inferCNV clusters with median values indicated (**a**) and for each spot on the Visium slide (**b**).

**Supplementary Fig. S16: Location of subclone differentially expressed genes mapped to selected subclone-specific CNAs in patient 1.** InferCNV signal is shown for selected chromosomes after averaging across all spots in each of the subclones. Vertical dashed lines indicate genomic positions of genes most highly expressed in the red subclone (defined as being differentially up-regulated in the red subclone in at least one of the pairwise comparisons and having the highest expression level in the red subclone when compared to all other subclones).

**Supplementary Fig. S17: Location of subclone differentially expressed genes mapped to selected subclone-specific CNAs in patient 2.** InferCNV signal is shown for selected chromosomes after averaging across all spots in each of the subclones. Vertical dashed lines indicate genomic positions of genes most highly expressed in the brown_red subclone (defined as being differentially up-regulated in the brown_red subclone in at least one of the pairwise comparisons and having the highest expression level in the brown_red subclone when compared to all other subclones).

**Supplementary Fig. S18: Location of subclone differentially expressed genes mapped to selected subclone-specific CNAs in patient 4.** InferCNV signal is shown for selected chromosomes after averaging across all spots in each of the subclones. Vertical dashed lines indicate genomic positions of genes most highly expressed in the brown subclone (defined as being differentially up-regulated in the brown subclone in at least one of the pairwise comparisons and having the highest expression level in the brown subclone when compared to all other subclones).

**Supplementary Fig. S19: Location of subclone differentially expressed genes mapped to selected subclone-specific CNAs in patient 5.** InferCNV signal is shown for selected chromosomes after averaging across all spots in each of the subclones. Vertical dashed lines indicate genomic positions of genes most highly expressed in the red (**a**) or blue (**b**) subclone (defined as being differentially up-regulated in the corresponding subclone in at least one of the pairwise comparisons and having the highest expression level in the corresponding subclone when compared to all other subclones).

**Supplementary Fig. S20: Location of subclone differentially expressed genes mapped to selected subclone-specific CNAs in patient 6.** InferCNV signal is shown for selected chromosomes after averaging across all spots in each of the subclones. Vertical dashed lines indicate genomic positions of genes most highly expressed in the red (**a**) or blue (**b**) subclone (defined as being differentially up-regulated in the corresponding subclone in at least one of the pairwise comparisons and having the highest expression level in the corresponding subclone when compared to all other subclones).

**Supplementary Fig. S21: Expression and detection rates of putative tumour intrinsic ligands in Zhang *et al*. scRNA-seq dataset** ^25^. Ligands identified in different samples are shown separately and are further split into ligands that were most highly expressed in tumour cells in our scRNA-seq dataset and ligand that were deemed plausibly subclone-specific as they were not detected in any of the cell types identified in our scRNA-seq dataset. Note that the latter did not show strong expression in any of the non-malignant subtypes. EOC, epithelial ovarian carcinoma.

**Supplementary Fig. S22: Giotto cell type enrichment in inferCNV clusters.** Dot colour shows PAGE enrichment scores averaged across all spots in each inferCNV cluster and each cell type; dot size shows percentage of spots in each inferCNV cluster that were significantly enriched for the corresponding cell type (significance determined using a threshold of 0.05 on Benjamini-Hochberg adjusted Giotto p-values). All malignant and non-malignant inferCNV clusters are shown for each sample, purple bars indicate the malignant clusters.

**Supplementary Fig. S23: Expression and detection rates of *CXCL10-CXCR3* and *CD47-SIRPA* ligand-receptor pairs in the Zhang *et al*. scRNA-seq dataset** ^25^.

**Supplementary Fig. S24: InferCNV heatmaps generated for Visium data on HGSOC samples from the Stur *et al*. study** ^63^. Heatmaps generated by inferCNV showing inferred CNA profiles of Visium spots are shown for samples from six excellent and six poor responders to neoadjuvant chemotherapy.

**Supplementary Fig. S25: Integrated gene expression analysis of eight Visium samples. a)** UMAP of eight integrated samples coloured by the sample of origin. **b)** UMAP of eight integrated samples coloured by the cluster. **c)** Top overexpressed genes in clusters 2 and 6 (candidate stromal dataset). **d)** Subclustering of candidate stromal dataset (clusters 2 and 6 from (**b**). **e)** Top overexpressed genes in cluster 3 of the candidate stromal dataset.

**Supplementary Fig. S26: Expression of marker genes used for Giotto cell type enrichment analysis in the scRNA-seq dataset.**

