## Supplementary Note 1 for "Spatial transcriptomics reveals ovarian cancer subclones with distinct tumour microenvironments"

### Supplementary Note 1. Good chemotherapy response is associated with stronger immune cell and fibroblast infiltration

To investigate possible differences between tumours of good and poor responders, we compared gene expression profiles of Visium spots annotated as malignant based on their CNA profiles in CRS1 and CRS3 patients. Differential gene expression analysis showed that in CRS3 (good responders) there were higher expression levels of genes involved in immune response (e.g. immunoglobulins and human leukocyte antigens) and extracellular matrix organisation (e.g. collagens, *DCN, FN1*) (**Supplementary Table ST10**). In contrast, genes previously implicated in ovarian cancer progression (*CLU, CD24, KRT7*)[1–5] and genes encoding ribosomal proteins were overexpressed in CRS1 (poor responder samples) (**Supplementary Table ST10**).

We examined the expression of these response-associated genes in our scRNA-seq dataset and found that genes over-expressed in the good responder samples were more specific to B cells, T cells, macrophages, and fibro5 (*FN1, COL3A1*) (**Figure SN1a**). In contrast, the genes over-expressed in the poor responder samples were more highly expressed in cancer cells or myofibroblasts (*ADIRF, MFGE8*) (**Figure SN1b**).

We used Bisque[6] to perform cell type decomposition of the malignant areas in CRS1 and CRS3 samples and found that, in agreement with the above, CRS3 patients have higher percentages of macrophages and fibro5 (*FN1, COL3A1*), whereas CRS1 patients have higher percentages of cancer cells, myofibroblasts, and mesothelial cells (**Figure SN1c**). We repeated decomposition analysis with CIBERSORTx[7] and it supported the conclusions (**Figure SN1d**). For both Bisque and CIBERSORTx, we used the full raw gene count matrix for our scRNA-seq dataset and pseudo-bulk counts for each of the six Visium samples (Visium spot counts summed up across malignant spots). Bisque was run with the default parameters. CIBERSORTx was run with S-mode for batch correction. Taken together, these results indicate stronger infiltration with immune cells and fibro5 in our good responder samples good_response_P2, good_response_P3, good_response_P5.

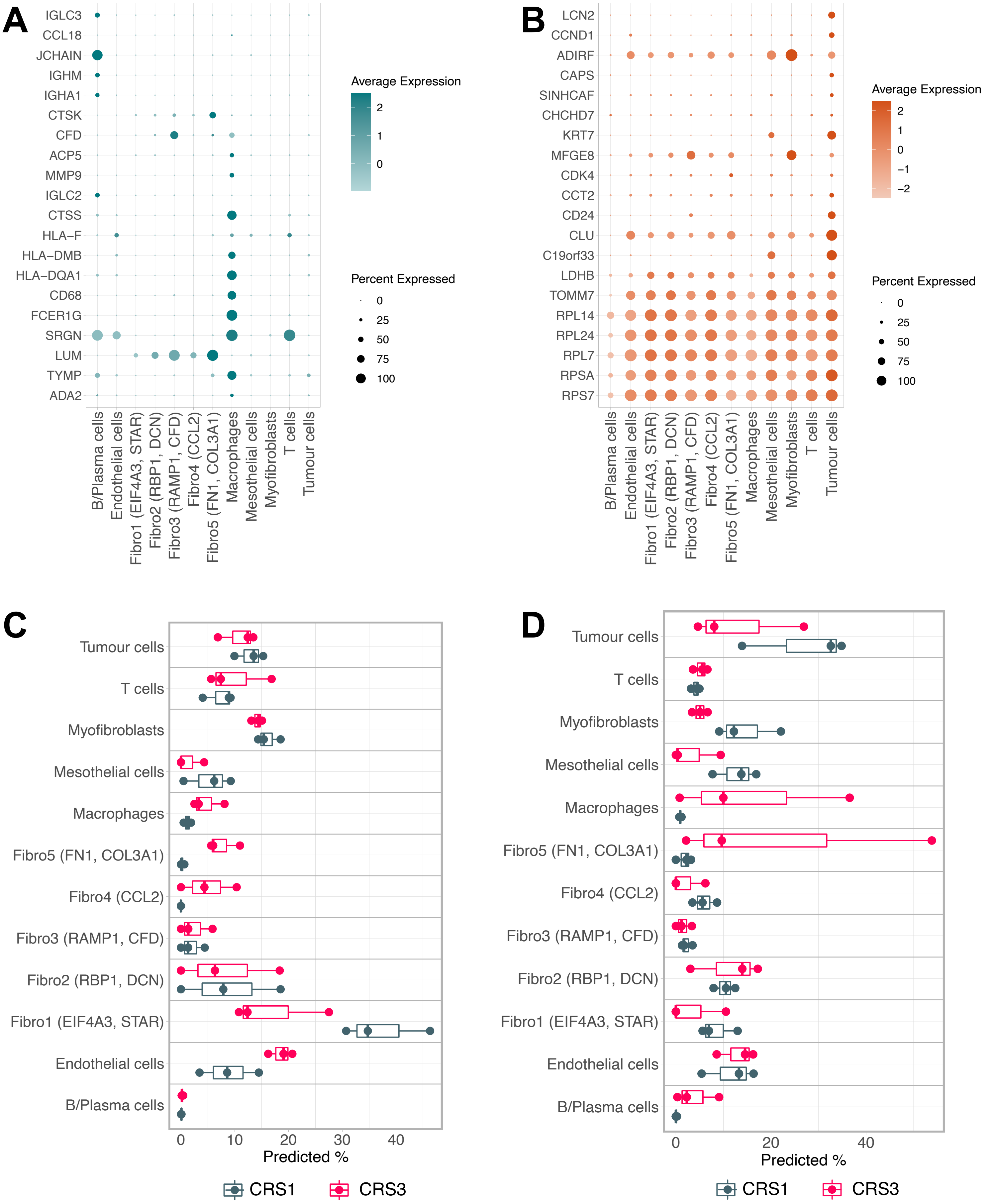


**a,b)** Expression in the scRNA-seq dataset of top 20 genes (based on the detection ratio) associated with **a**) good response to chemotherapy. **b)** poor response to chemotherapy. **c)** Cell type deconvolution of malignant tissue areas in good (CRS3, shown in red) and poor responders (CRS1, shown in grey) using Bisque. **d)** Cell type deconvolution of malignant areas in good (CRS3, shown in red) and poor (CRS1, shown in grey) responders using CIBERSORTx.
