## Supplementary Figures for "Spatial transcriptomics reveals ovarian cancer subclones with distinct tumour microenvironments"

### Fig. S1

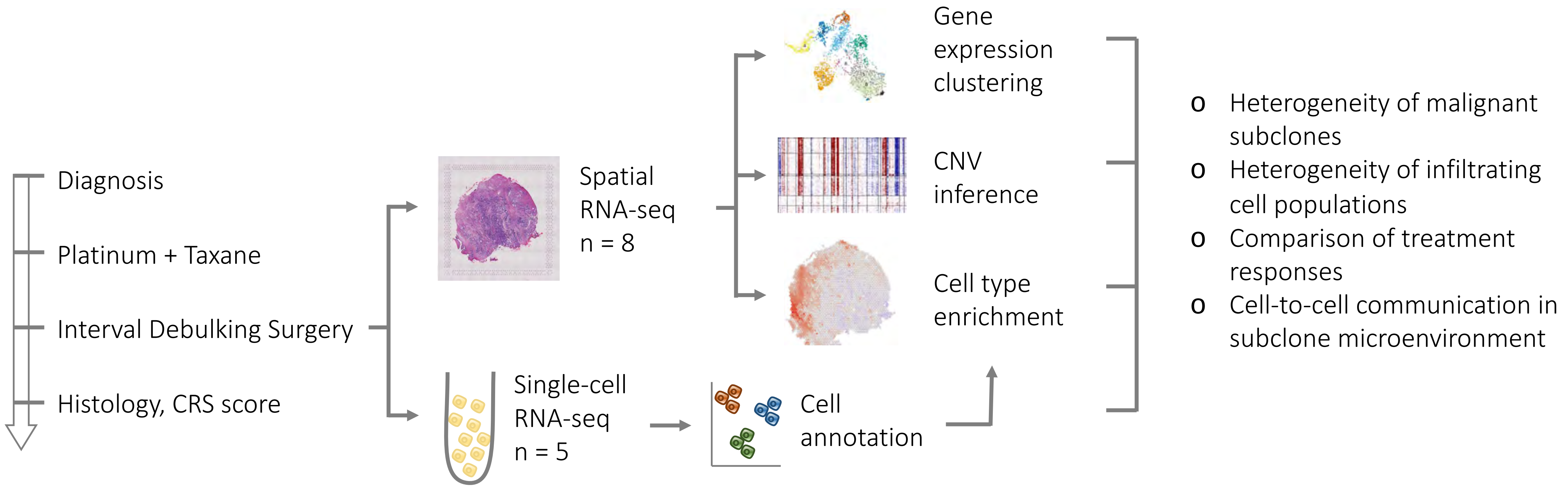


**Study design.** Samples were collected during interval debulking surgery from patients who underwent 3-4 cycles of platinum and taxane treatment. Eight samples were profiled using 10x Genomics Visium and five samples using 10x Genomics 3' Gene Expression solution.

### Fig. S2

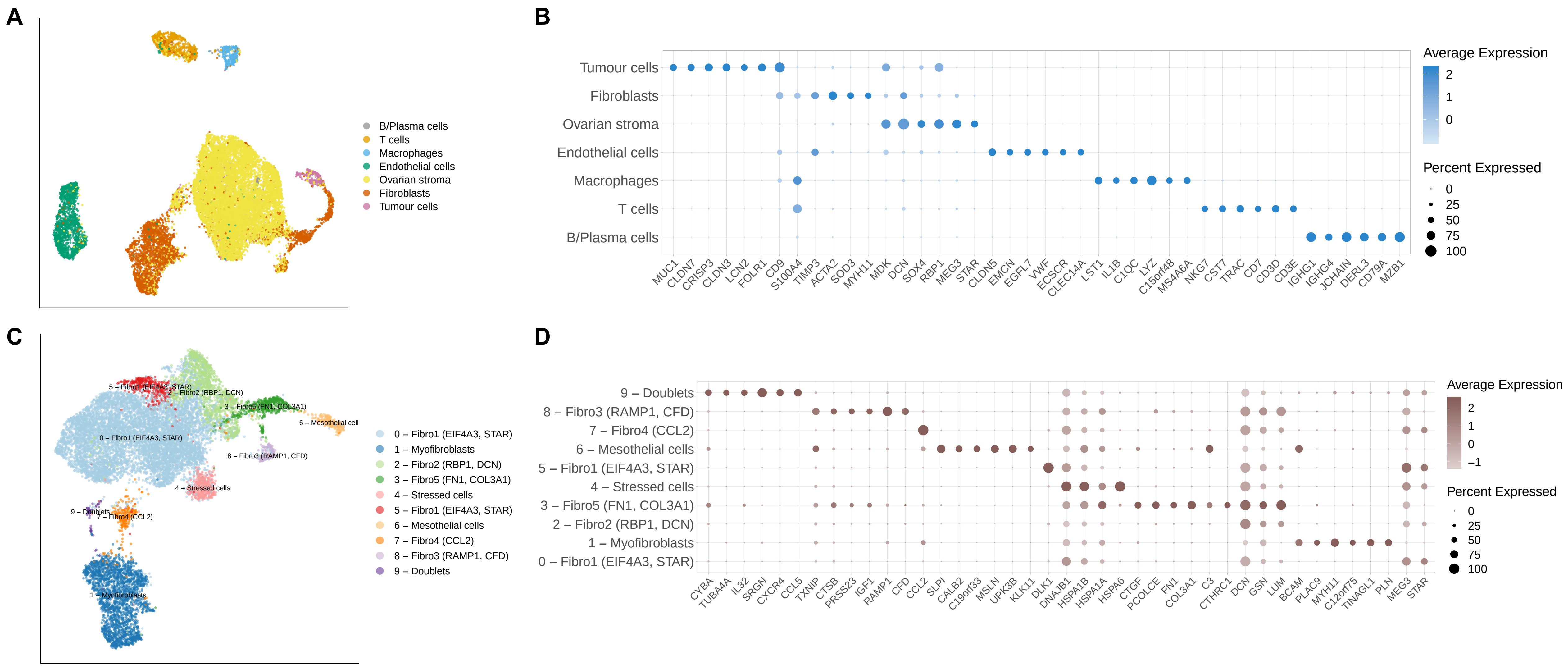


**Annotation of the single-cell RNA-seq HGSOC dataset. a)** UMAP showing top-level cell type annotations inferred by scMatch with Olbrecht *et al.* reference dataset. **b)** Differentially expressed genes (DEGs) identified in the top-level cell types; top DEGs based on the ratio of detection rates are shown for each cell type. **c)** UMAP showing subclustering of fibroblasts and stromal cells. **d)** DEGs identified in the subclusters shown in (**c**); top DEGs based on the ratio of detection rates are shown for each cell type.

### Fig. S3

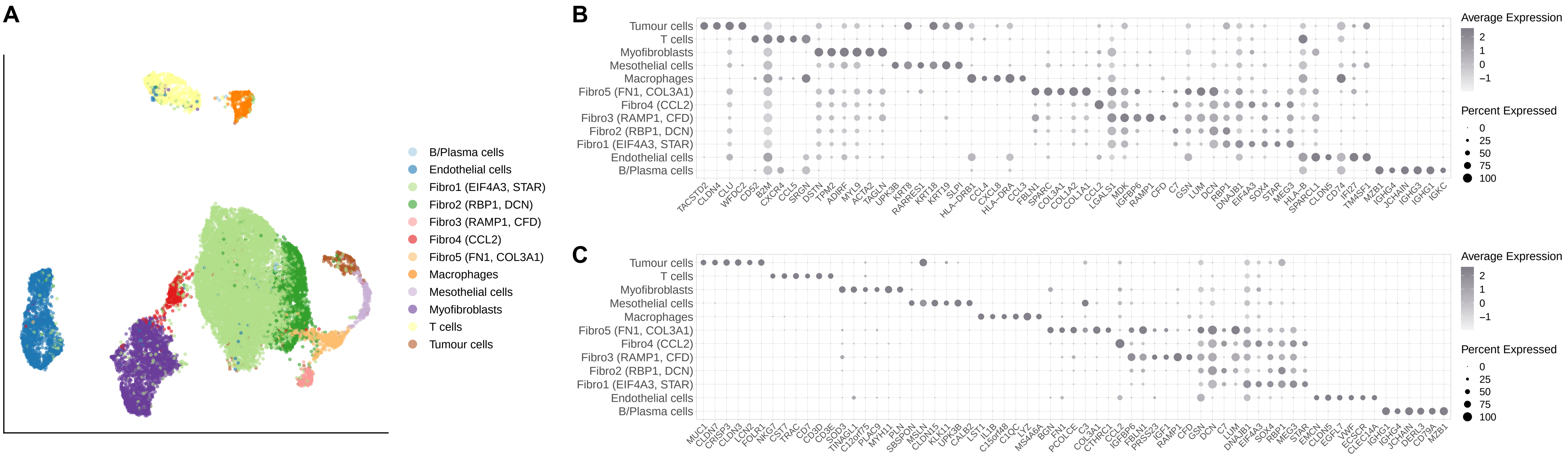


**Fine-grain annotations of the scRNA-seq dataset. a)** UMAP showing 12 cell types. **b)** Top differentially expressed genes (DEGs) over-expressed in each of the cell types, ranked based on logFC. **c)** Top DEGs over-expressed in each of the cell types, ranked based on the ratio of detection rates. DEGs were calculated for each cell type vs all other cells.

### Fig. S4

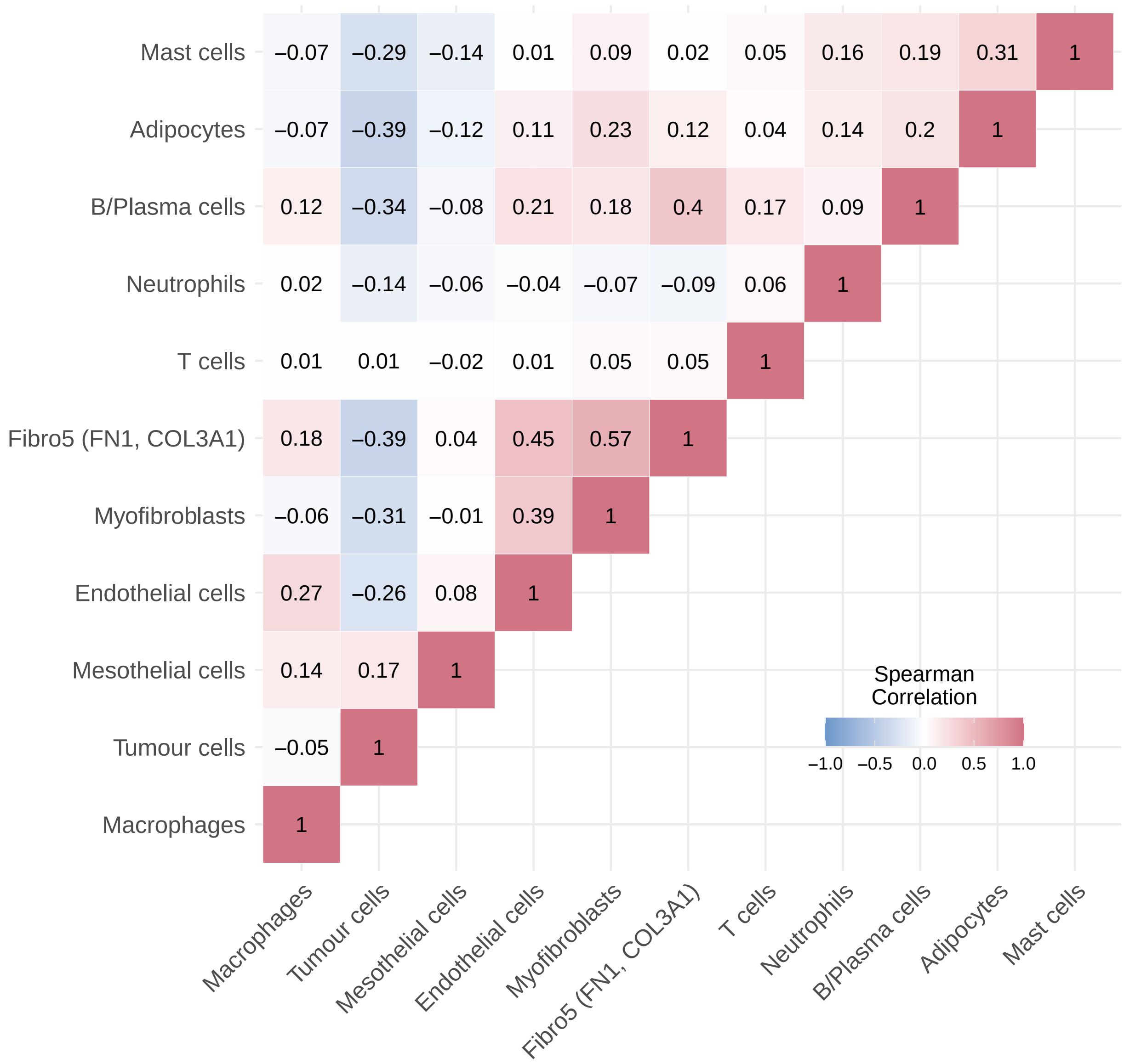


**Correlation between Giotto cell type enrichment scores across all Visium samples and spots.**

### Fig. S5

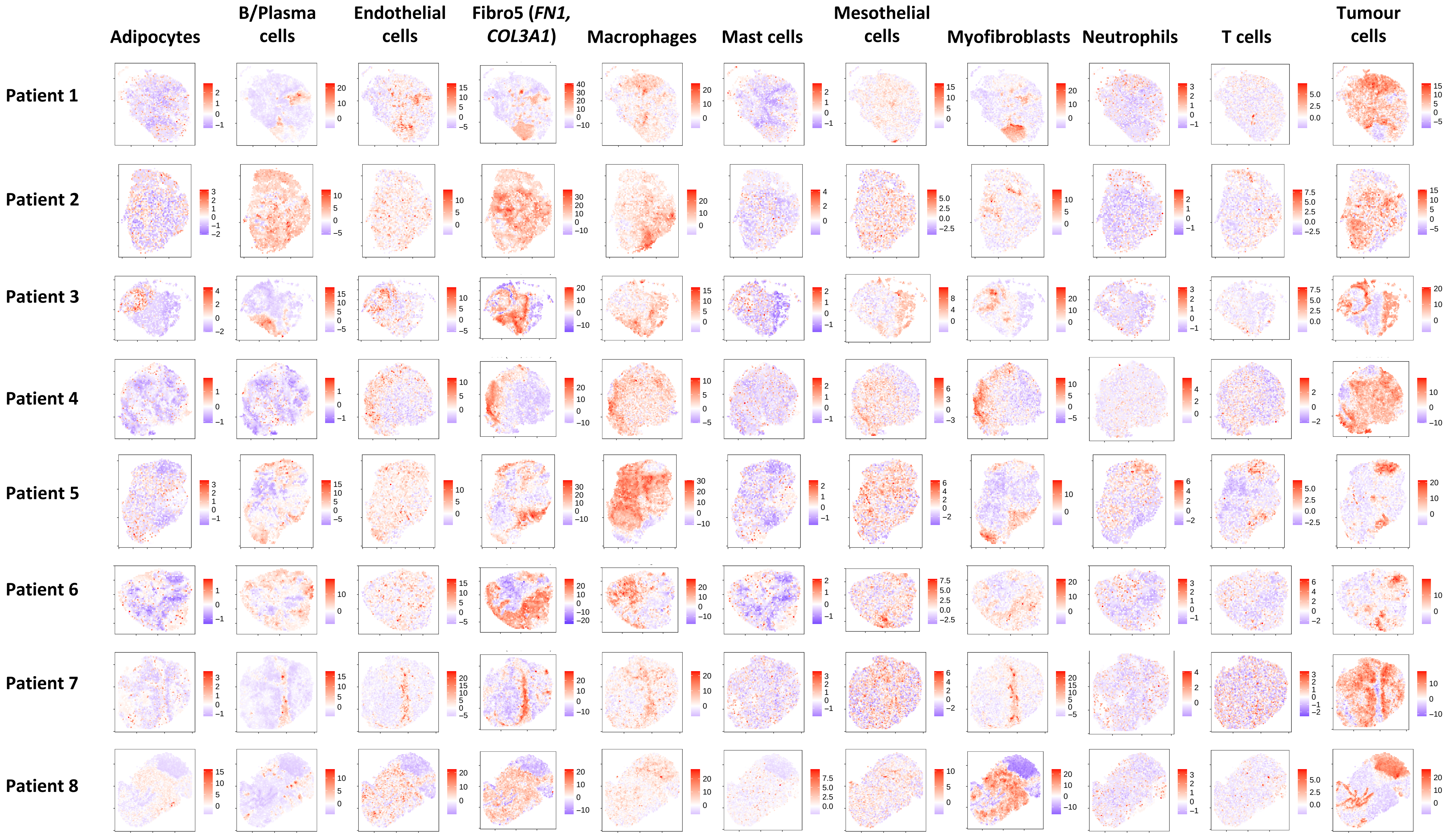


**Giotto cell type enrichment scores for eleven cell populations in Visium samples.**

Fig. S6

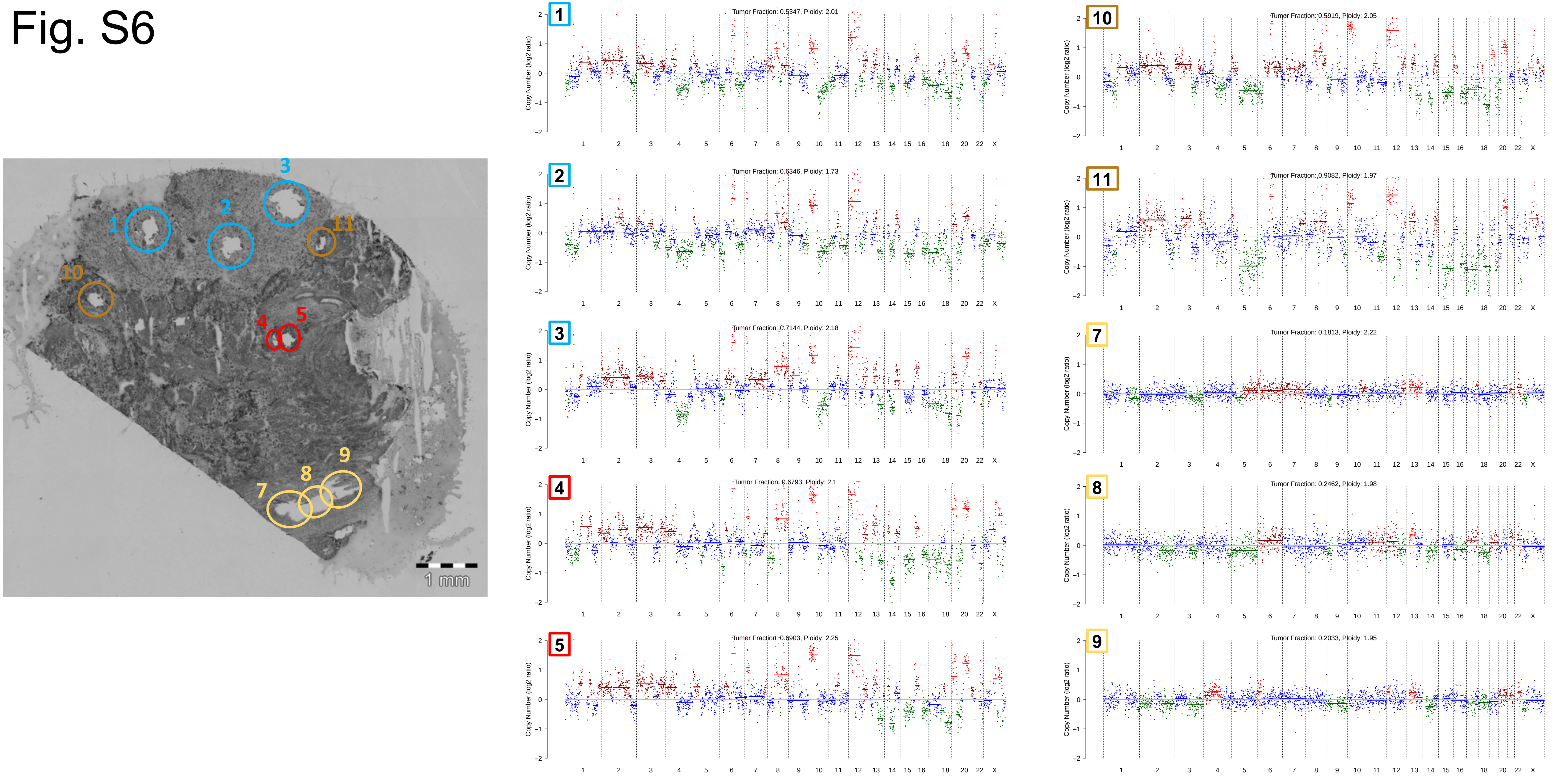


**Validation of inferCNV predictions using ultra-low-pass whole genome sequencing.** IchorCNV analysis was performed for each of the numbered regions separately, using tongue tissue as a background. IchorCNV CNA profiles are shown for each region, where green indicates 1 copy, blue 2 copies, brown 3 copies, red 4+ copies.

### Fig. S7

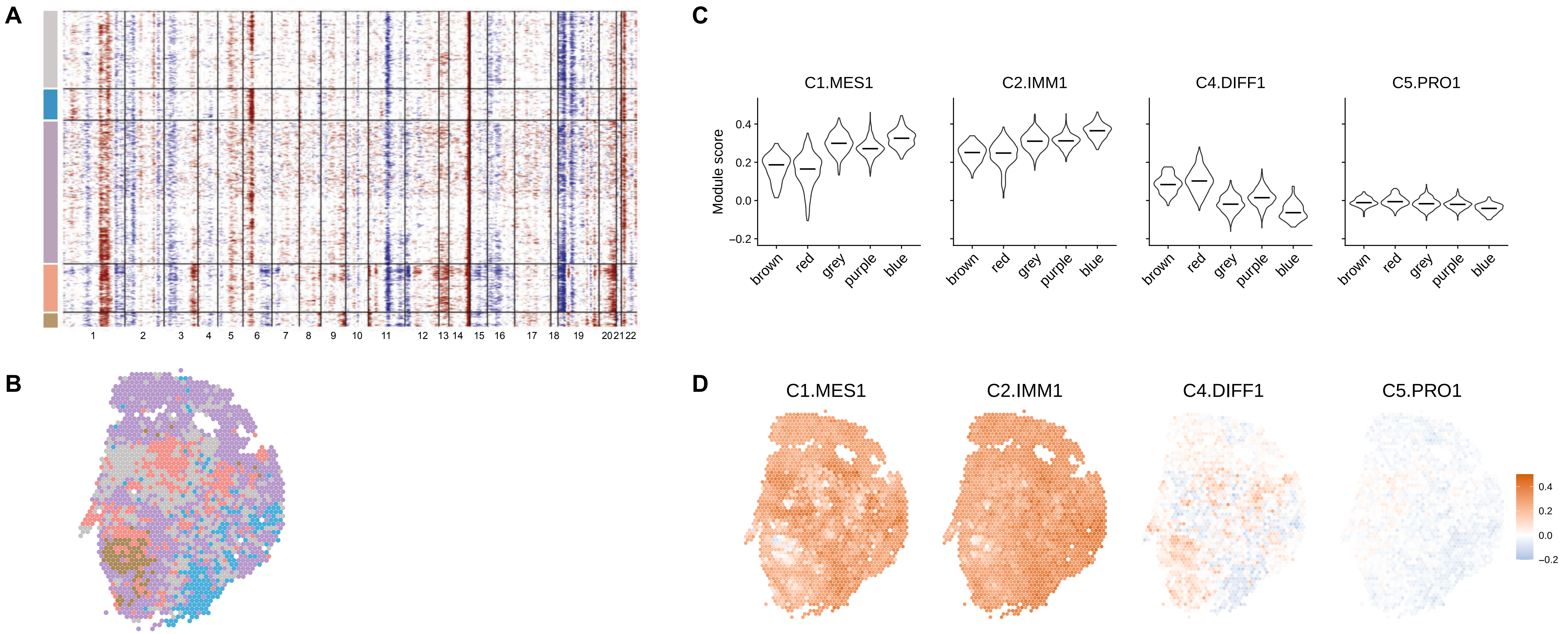


**Visium data summary for patient 2**. **a)** Heatmap generated by inferCNV showing inferred CNA profiles of Visium spots. Horizontal black lines separate clusters identified by inferCNV. Red corresponds to predicted amplification, blue to predicted deletion. **b)** Projection of spot clusters identified by inferCNV onto the tissue section. **c, d)** Module scores calculated using *AddModuleScore* function in Seurat for genes overexpressed in four HGSOC molecular subtypes; shown in malignant inferCNV clusters with median values indicated (**c**) and for each spot on the Visium slide (**d**).

### Fig. S8

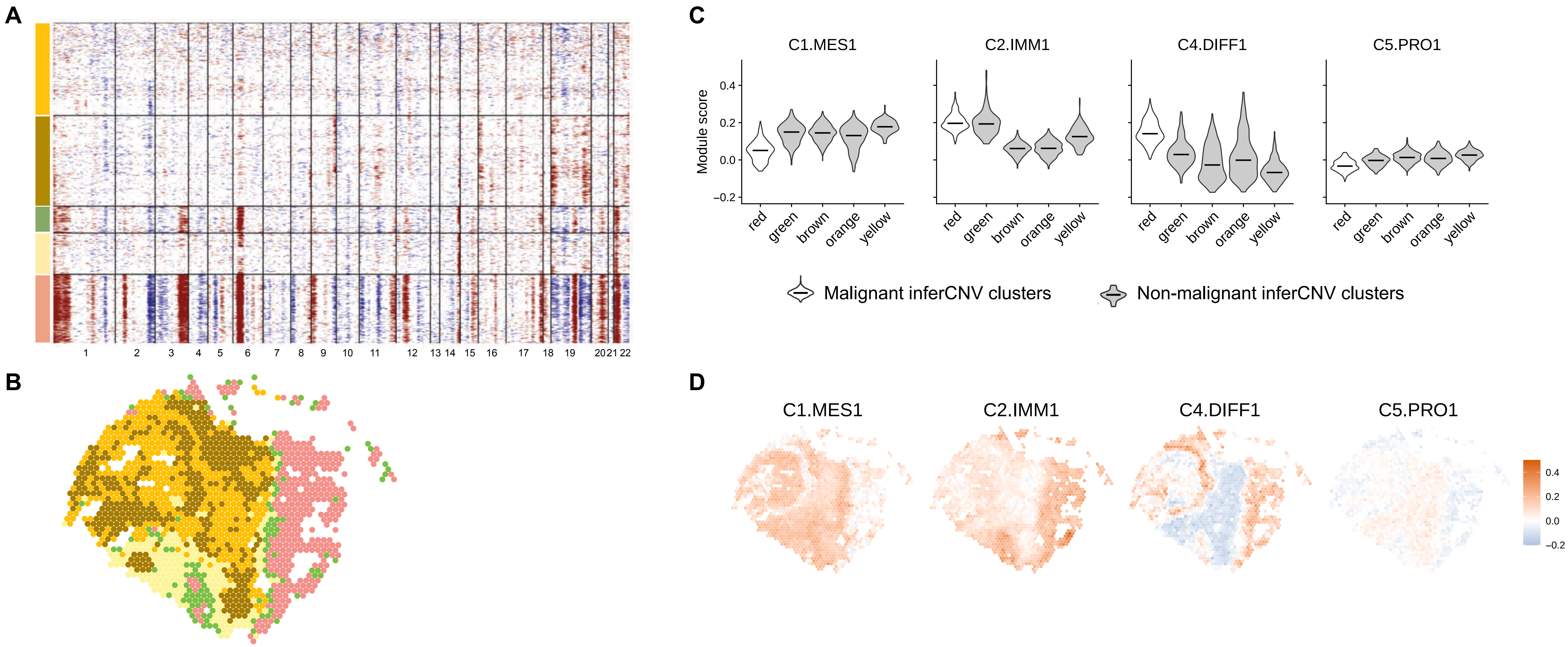


**Visium data summary for patient 3**. **a)** Heatmap generated by inferCNV showing inferred CNA profiles of Visium spots. Horizontal black lines separate clusters identified by inferCNV. Red corresponds to predicted amplification, blue to predicted deletion. **b)** Projection of spot clusters identified by inferCNV onto the tissue section. **c, d)** Module scores calculated using AddModuleScore function in Seurat for genes overexpressed in four HGSOC molecular subtypes; shown in malignant inferCNV clusters with median values indicated (**c**) and for each spot on the Visium slide (**d**).

### Fig. S9

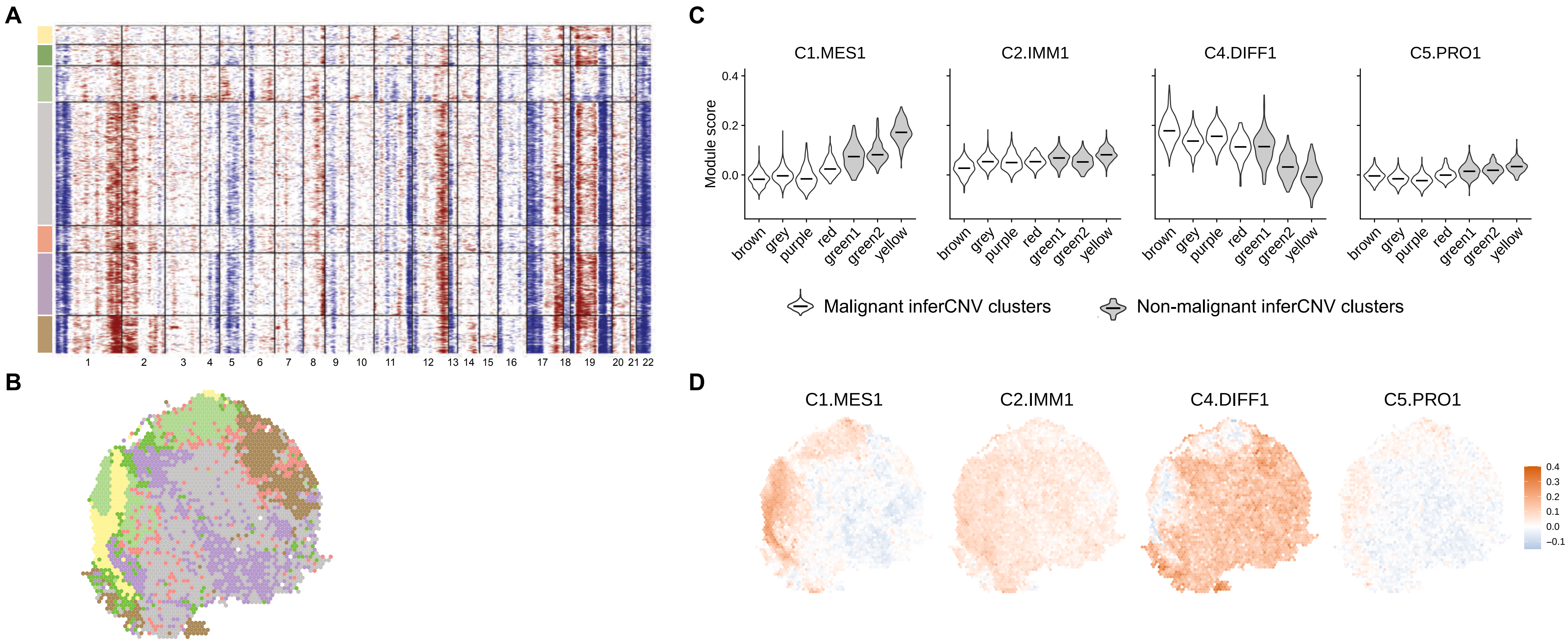


**Visium data summary for patient 4**. **a)** Heatmap generated by inferCNV showing inferred CNA profiles of Visium spots. Horizontal black lines separate clusters identified by inferCNV. Red corresponds to predicted amplification, blue – predicted deletion. **b)** Projection of spot clusters identified by inferCNV onto the tissue section. **c, d)** Module scores calculated using AddModuleScore function in Seurat for genes overexpressed in four HGSOC molecular subtypes; shown in malignant inferCNV clusters with median values indicated (**c**) and for each spot on the Visium slide (**d**).

### Fig. S10

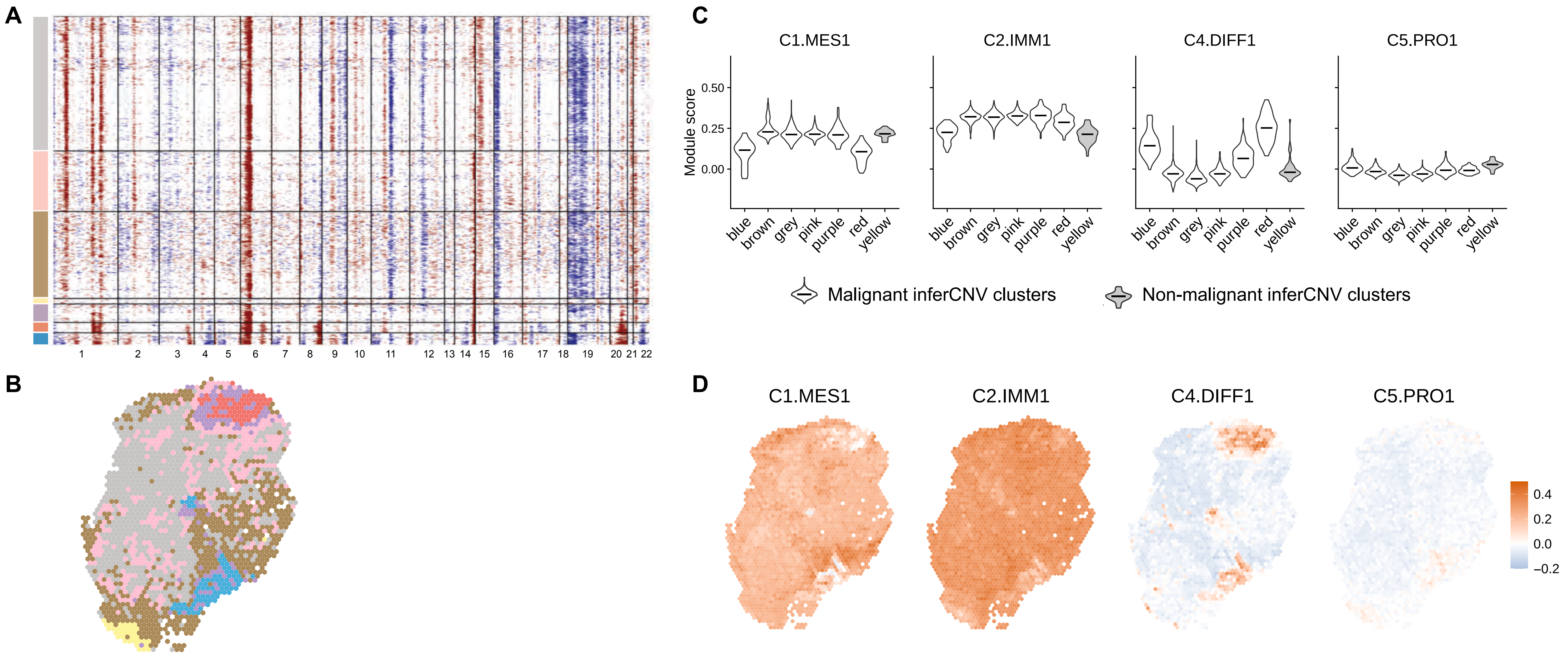


**Visium data summary for patient 5**. **a)** Heatmap generated by inferCNV showing inferred CNA profiles of Visium spots. Horizontal black lines separate clusters identified by inferCNV. Red corresponds to predicted amplification, blue – predicted deletion. **b)** Projection of spot clusters identified by inferCNV onto the tissue section. **c, d)** Module scores calculated using AddModuleScore function in Seurat for genes overexpressed in four HGSOC molecular subtypes; shown in inferCNV clusters with median values indicated (**c**) and for each spot on the Visium slide (**d**). Note, panels **b**, **c**, **d** are duplicating Figure 3.

### Fig. S11

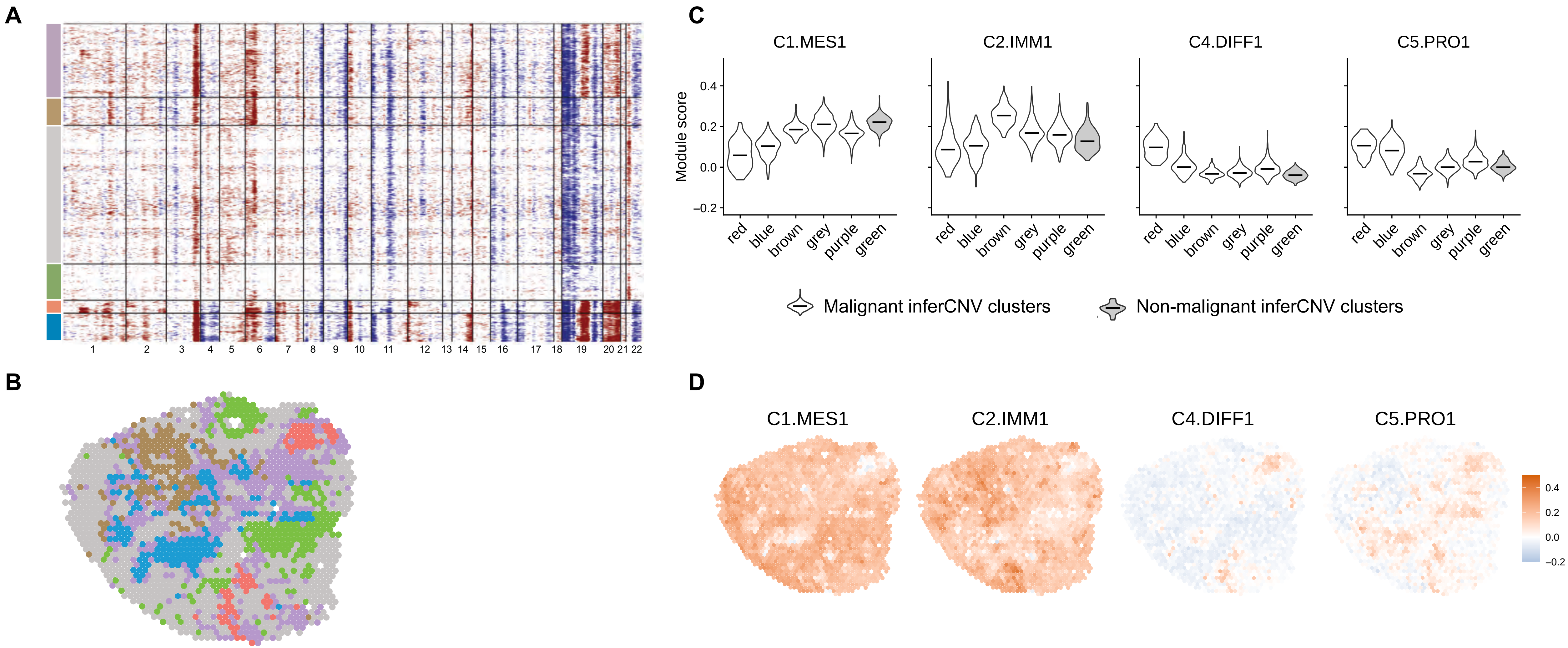


**Visium data summary for patient 6**. **a)** Heatmap generated by inferCNV showing inferred CNA profiles of Visium spots. Horizontal black lines separate clusters identified by inferCNV. Red corresponds to predicted amplification, blue – predicted deletion. **b)** Projection of spot clusters identified by inferCNV onto the tissue section. **c, d)** Module scores calculated using AddModuleScore function in Seurat for genes overexpressed in four HGSOC molecular subtypes; shown in malignant inferCNV clusters with median values indicated (**c**) and for each spot on the Visium slide (**d**).

### Fig. S12

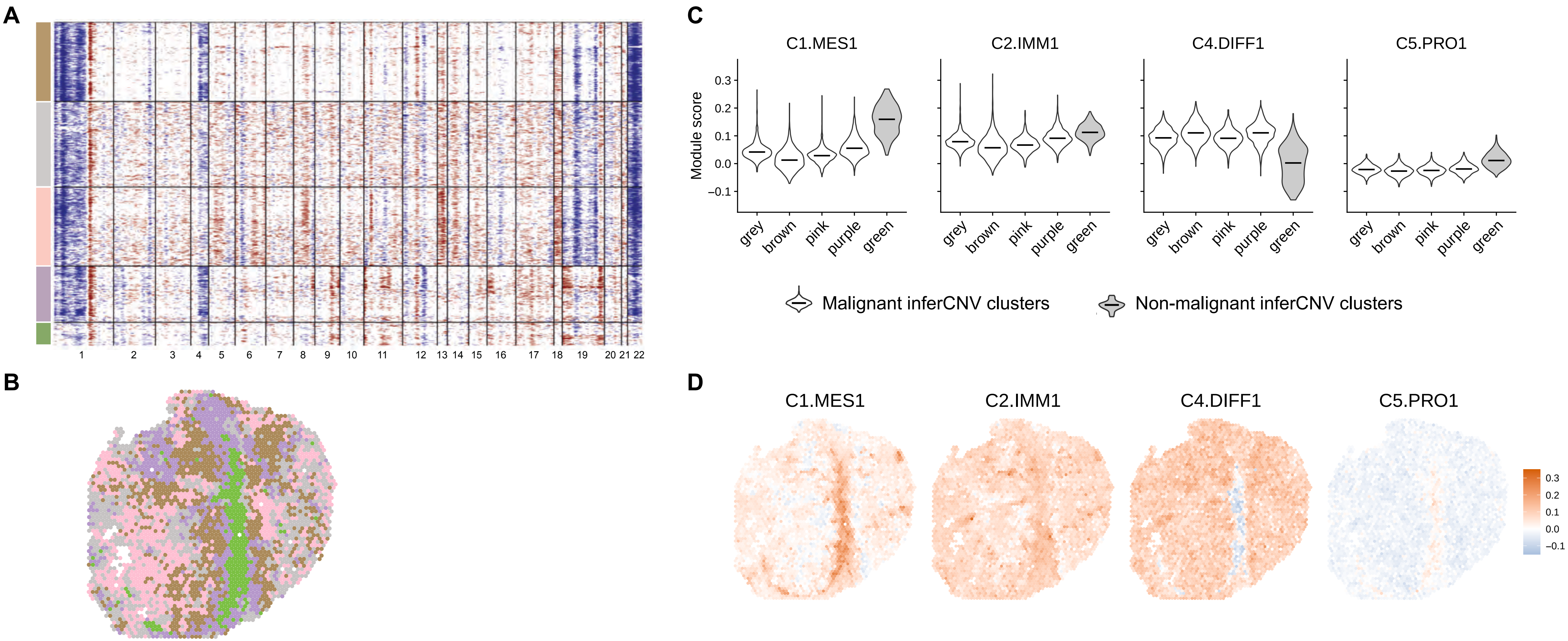


**Visium data summary for patient 7**. **a)** Heatmap generated by inferCNV showing inferred CNA profiles of Visium spots. Horizontal black lines separate clusters identified by inferCNV. Red corresponds to predicted amplification, blue – predicted deletion. **b)** Projection of spot clusters identified by inferCNV onto the tissue section. **c, d)** Module scores calculated using AddModuleScore function in Seurat for genes overexpressed in four HGSOC molecular subtypes; shown in malignant inferCNV clusters with median values indicated (**c**) and for each spot on the Visium slide (**d**).

### Fig. S13

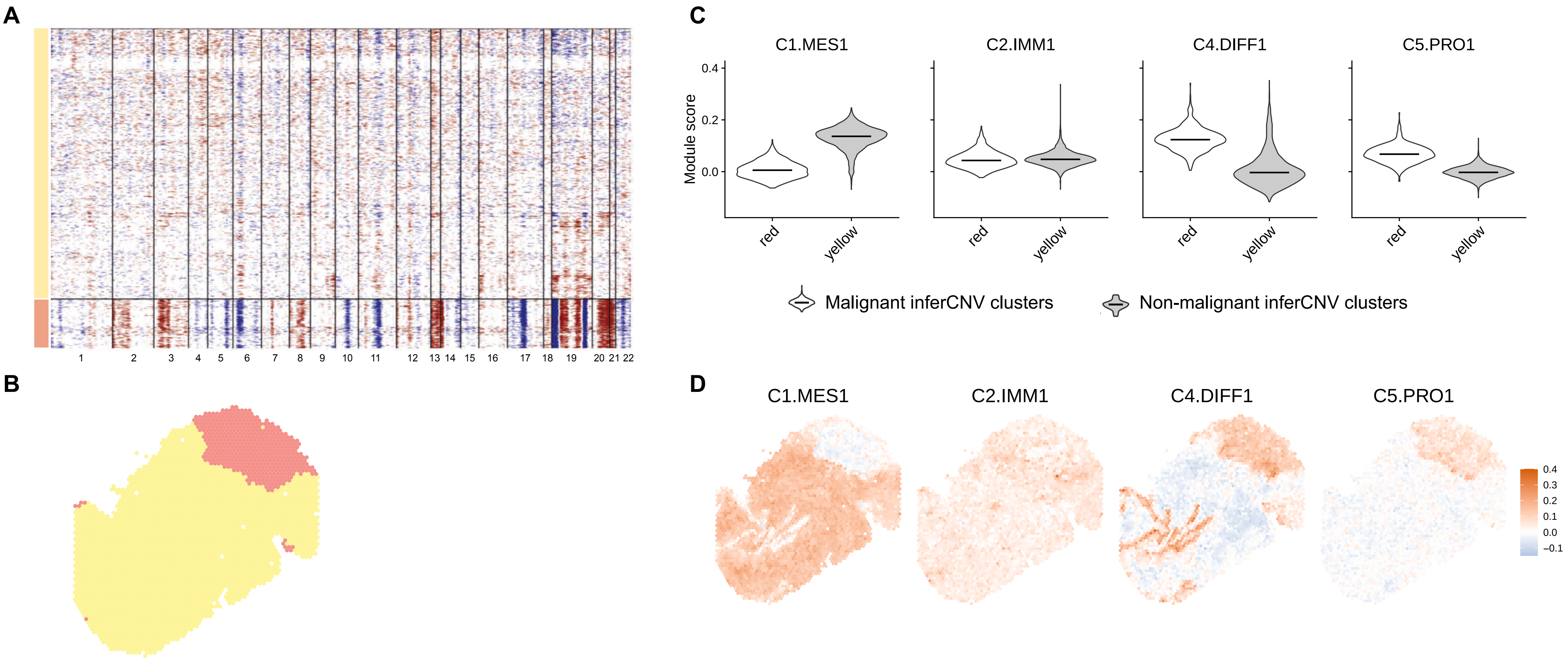


**Visium data summary for patient 8**. **a)** Heatmap generated by inferCNV showing inferred CNA profiles of Visium spots. Horizontal black lines separate clusters identified by inferCNV. Red corresponds to predicted amplification, blue – predicted deletion. **b)** Projection of spot clusters identified by inferCNV onto the tissue section. **c, d)** Module scores calculated using AddModuleScore function in Seurat for genes overexpressed in four HGSOC molecular subtypes; shown in malignant inferCNV clusters with median values indicated (**c**) and for each spot on the Visium slide (**d**).

### Fig. S14

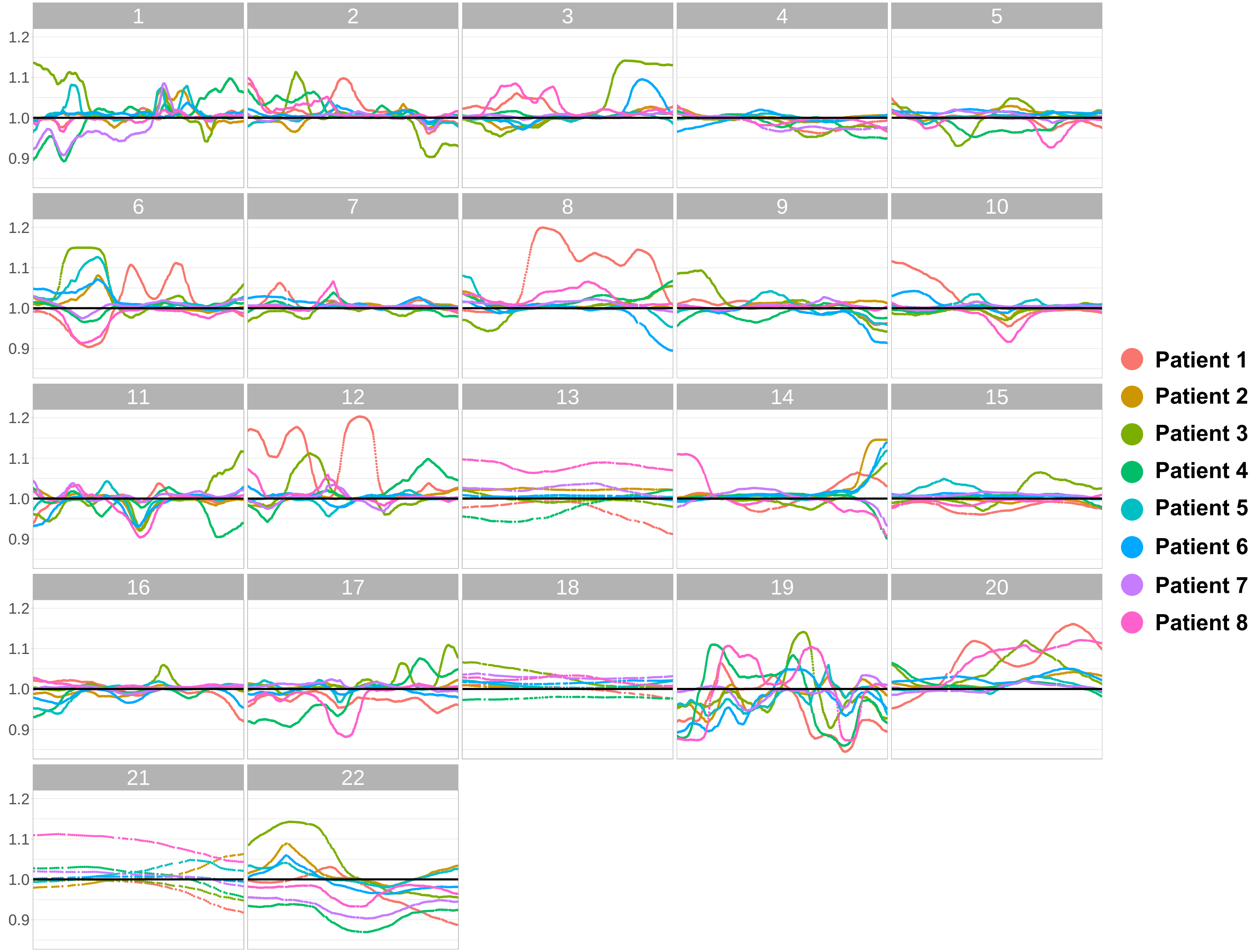


**Summary of inferCNV signal predicted in eight Visium samples**. Each panel shows one chromosome. InferCNV signal was averaged for each sample across all malignant spots. Genomic areas with InferCNV signal above 1.1 were considered amplified for the between-sample and TCGA dataset comparisons.

### Fig. S15

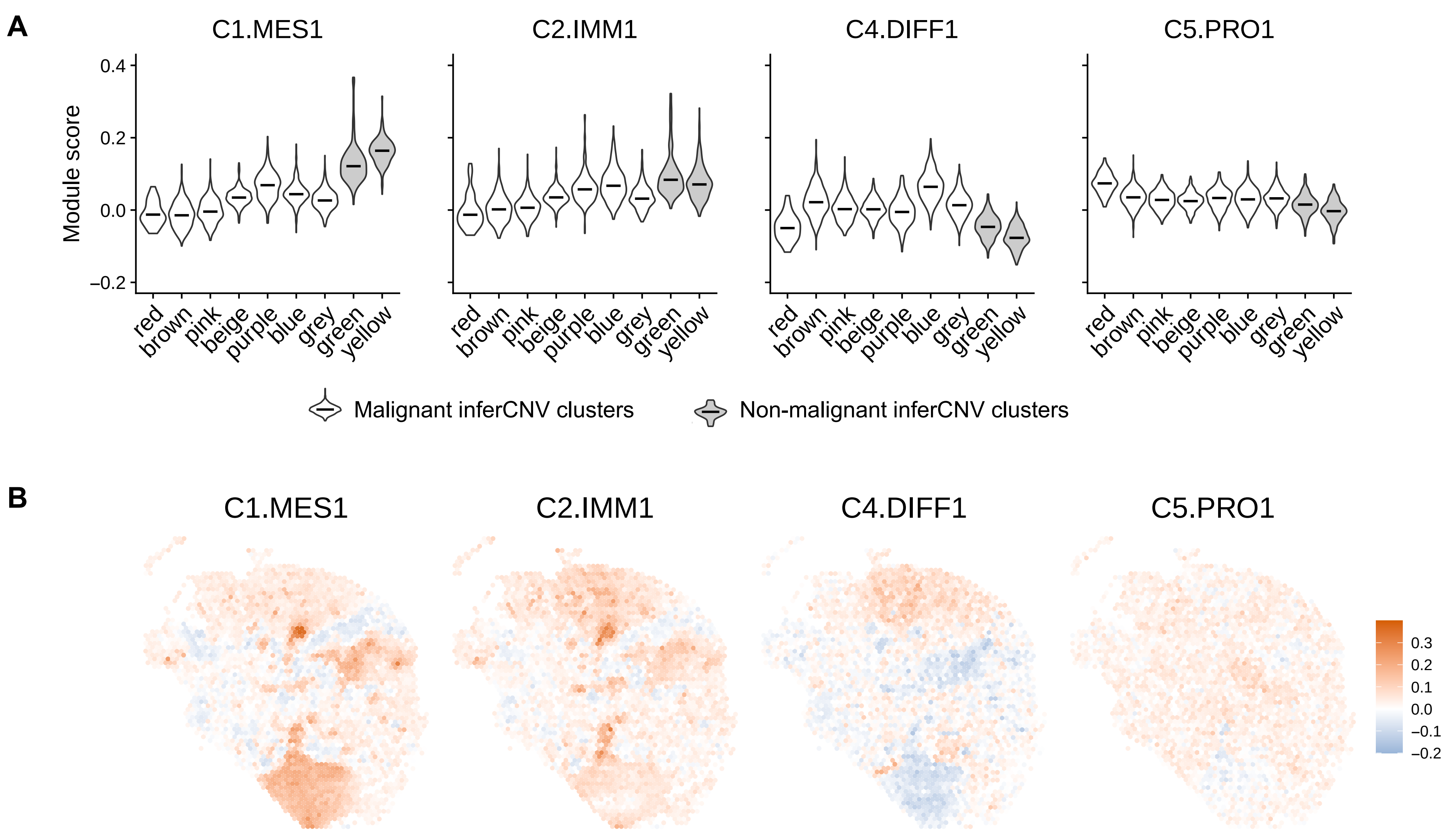


**Visium data summary for patient 1**. Module scores calculated using *AddModuleScore* function in Seurat for genes overexpressed in four HGSOC molecular subtypes; shown in malignant inferCNV clusters with median values indicated (**a**) and for each spot on the Visium slide (**b**).

### Fig. S16

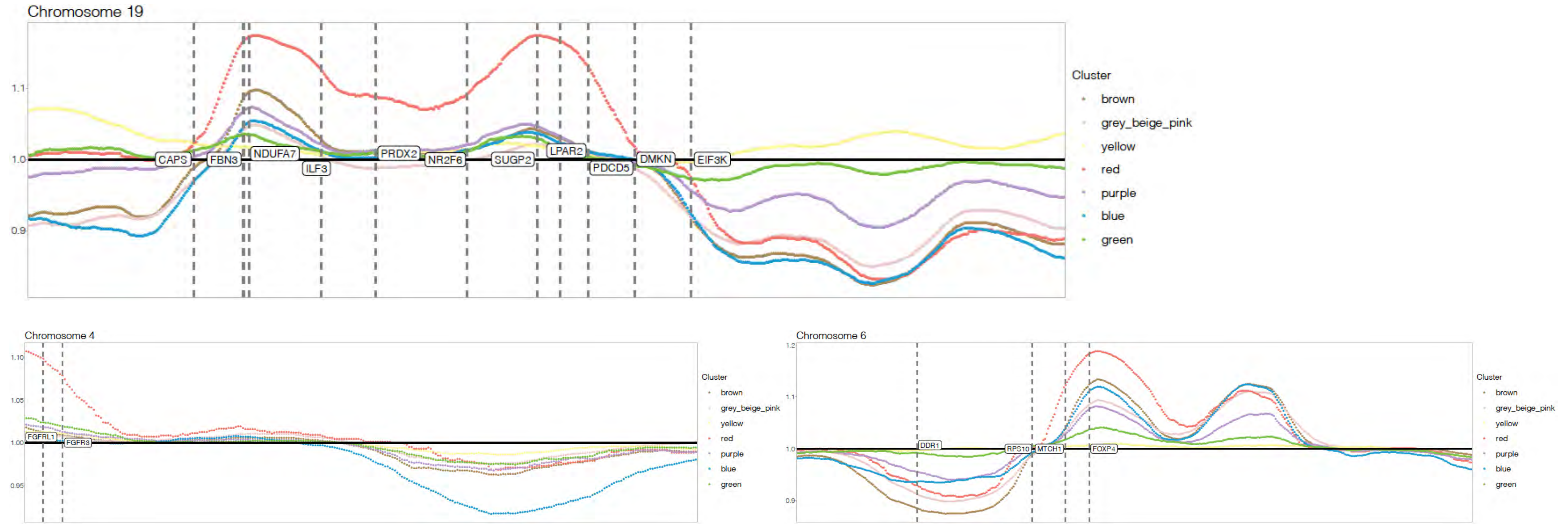


**Location of subclone differentially expressed genes mapped to selected subclone-specific CNAs in patient 1.** InferCNV signal is shown for selected chromosomes after averaging across all spots in each of the subclones. Vertical dashed lines indicate genomic positions of genes most highly expressed in the red subclone (defined as being differentially up-regulated in the red subclone in at least one of the pairwise comparisons and having the highest expression level in the red subclone when compared to all other subclones).

### Fig. S17

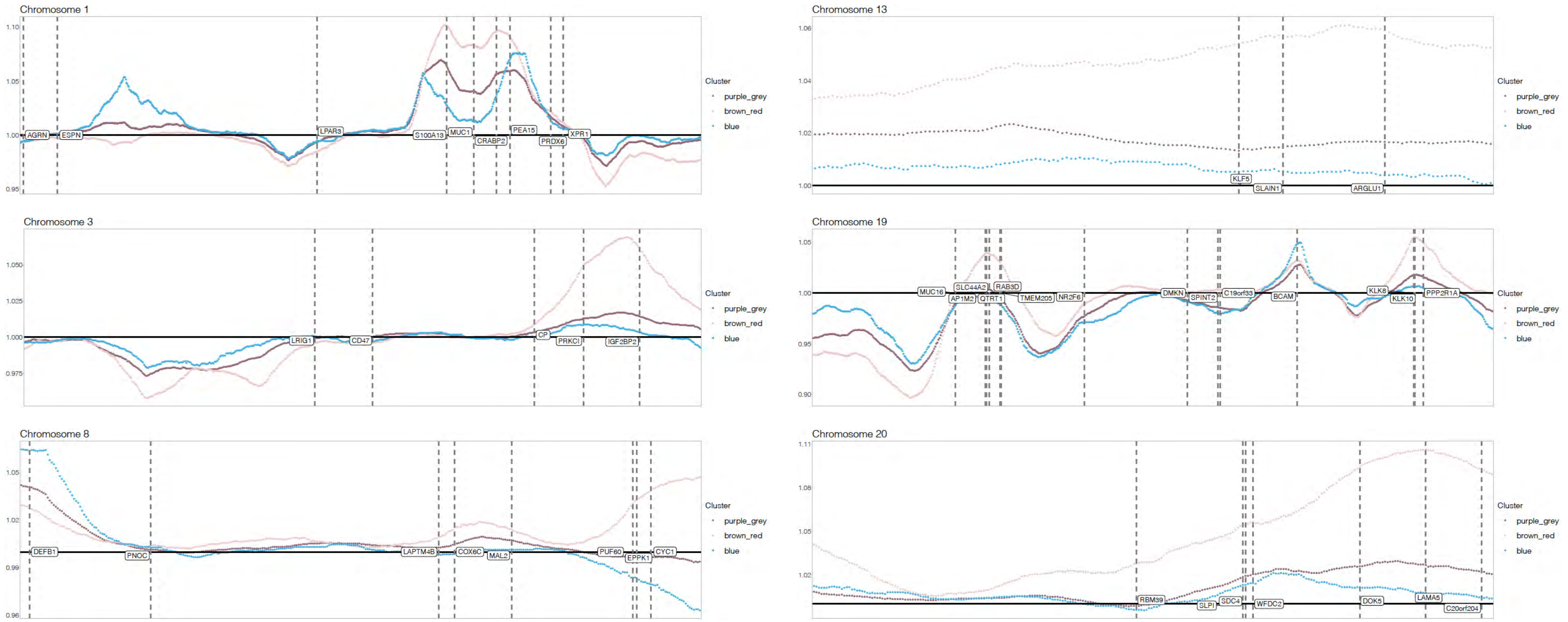


**Location of subclone differentially expressed genes mapped to selected subclone-specific CNAs in patient 2.** InferCNV signal is shown for selected chromosomes after averaging across all spots in each of the subclones. Vertical dashed lines indicate genomic positions of genes most highly expressed in the brown_red subclone (defined as being differentially up-regulated in the brown_red subclone in at least one of the pairwise comparisons and having the highest expression level in the brown_red subclone when compared to all other subclones).

### Fig. S18

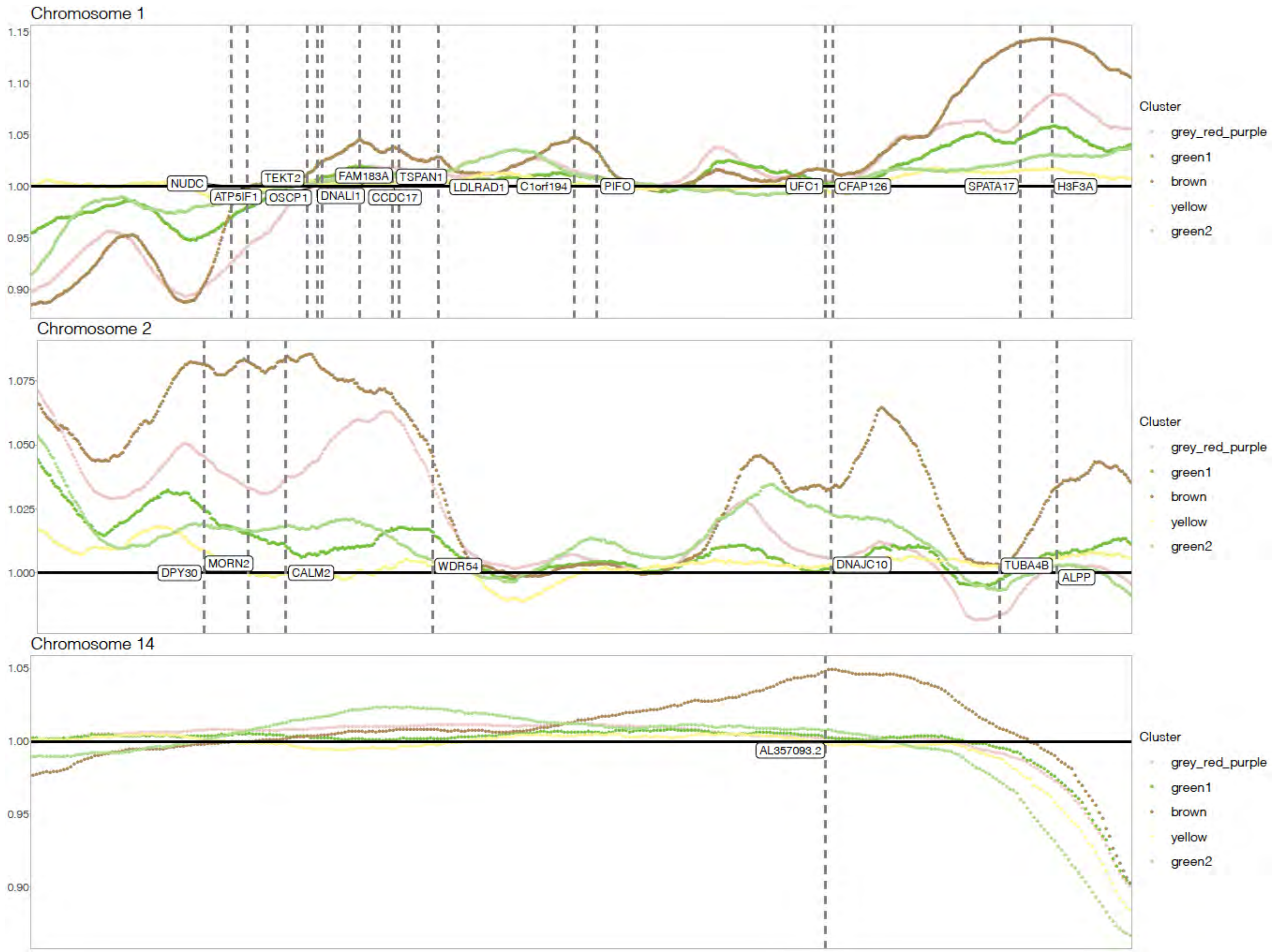


**Location of subclone differentially expressed genes mapped to selected subclone-specific CNAs in patient 4.** InferCNV signal is shown for selected chromosomes after averaging across all spots in each of the subclones. Vertical dashed lines indicate genomic positions of genes most highly expressed in the brown subclone (defined as being differentially up-regulated in the brown subclone in at least one of the pairwise comparisons and having the highest expression level in the brown subclone when compared to all other subclones).

### Fig. S19

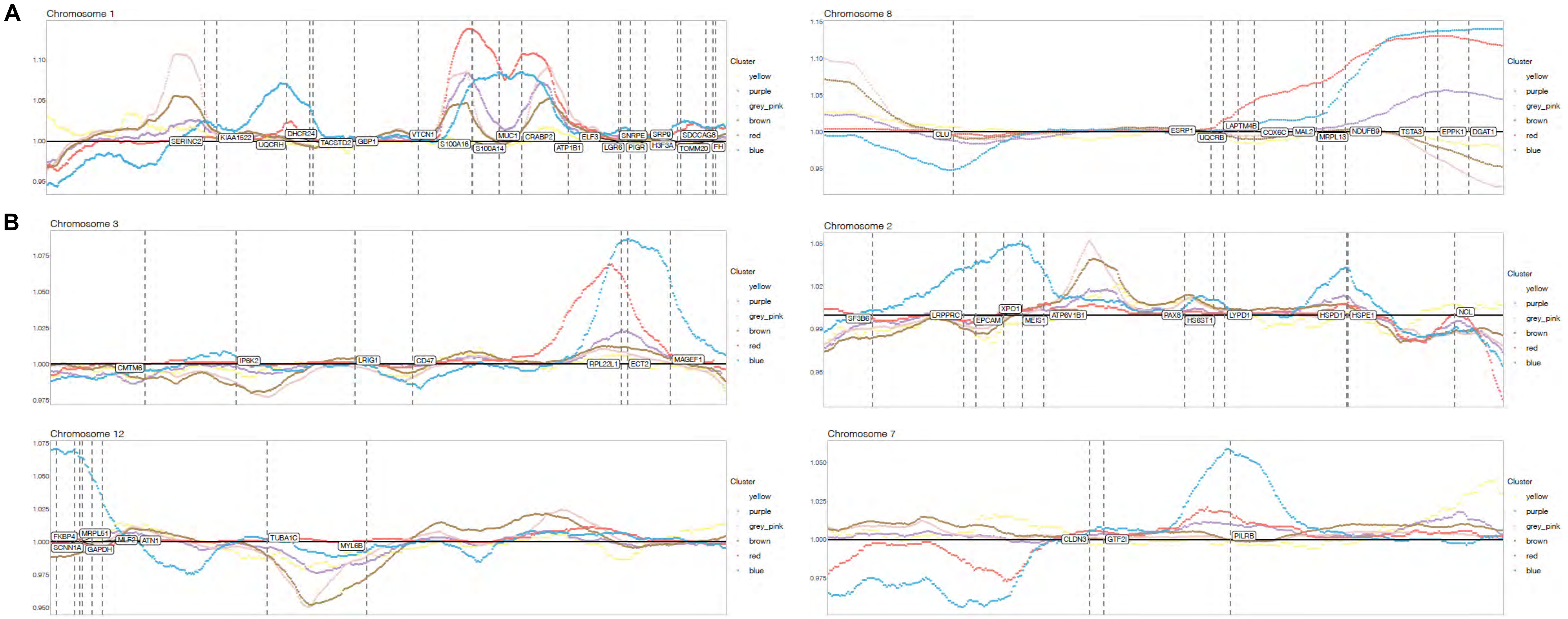


**Location of subclone differentially expressed genes mapped to selected subclone-specific CNAs in patient 5.** InferCNV signal is shown for selected chromosomes after averaging across all spots in each of the subclones. Vertical dashed lines indicate genomic positions of genes most highly expressed in the red (**a**) or blue (**b**) subclone (defined as being differentially up-regulated in the corresponding subclone in at least one of the pairwise comparisons and having the highest expression level in the corresponding subclone when compared to all other subclones).

### Fig. S20

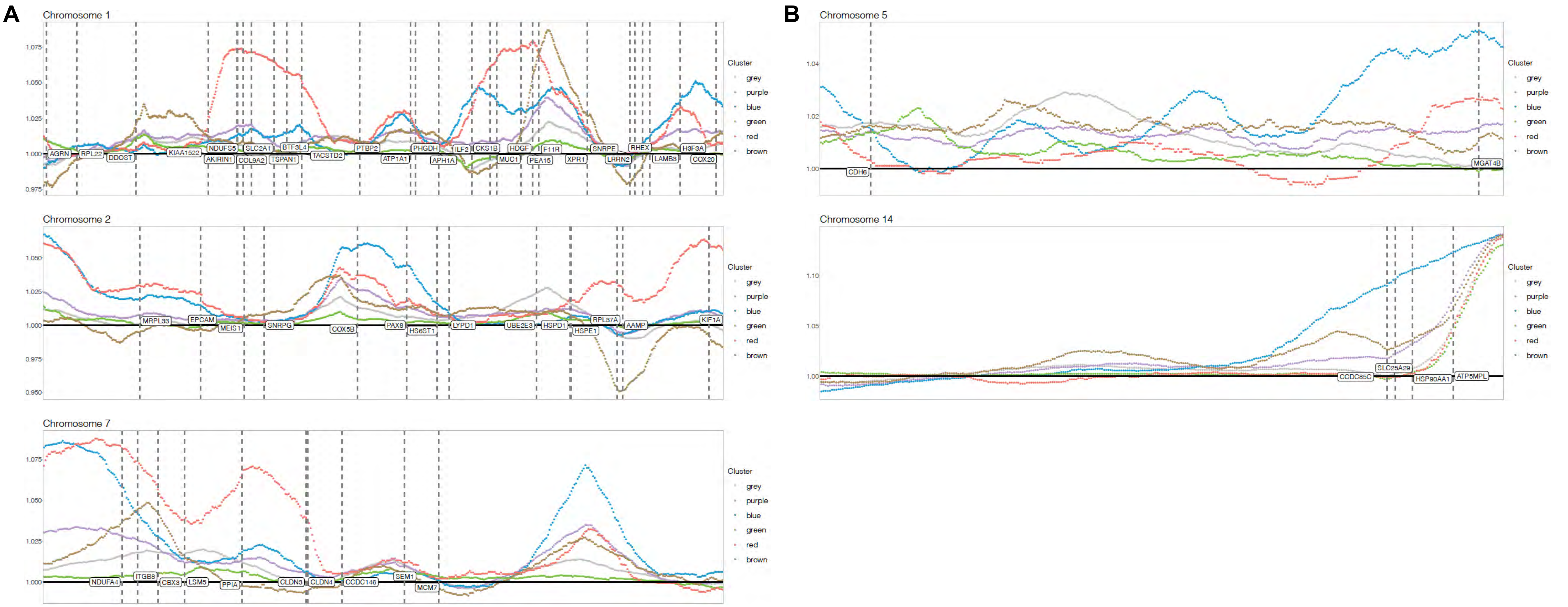


**Location of subclone differentially expressed genes mapped to selected subclone-specific CNAs in patient 6.** InferCNV signal is shown for selected chromosomes after averaging across all spots in each of the subclones. Vertical dashed lines indicate genomic positions of genes most highly expressed in the red (**a**) or blue (**b**) subclone (defined as being differentially up-regulated in the corresponding subclone in at least one of the pairwise comparisons and having the highest expression level in the corresponding subclone when compared to all other subclones).

### Fig. S21

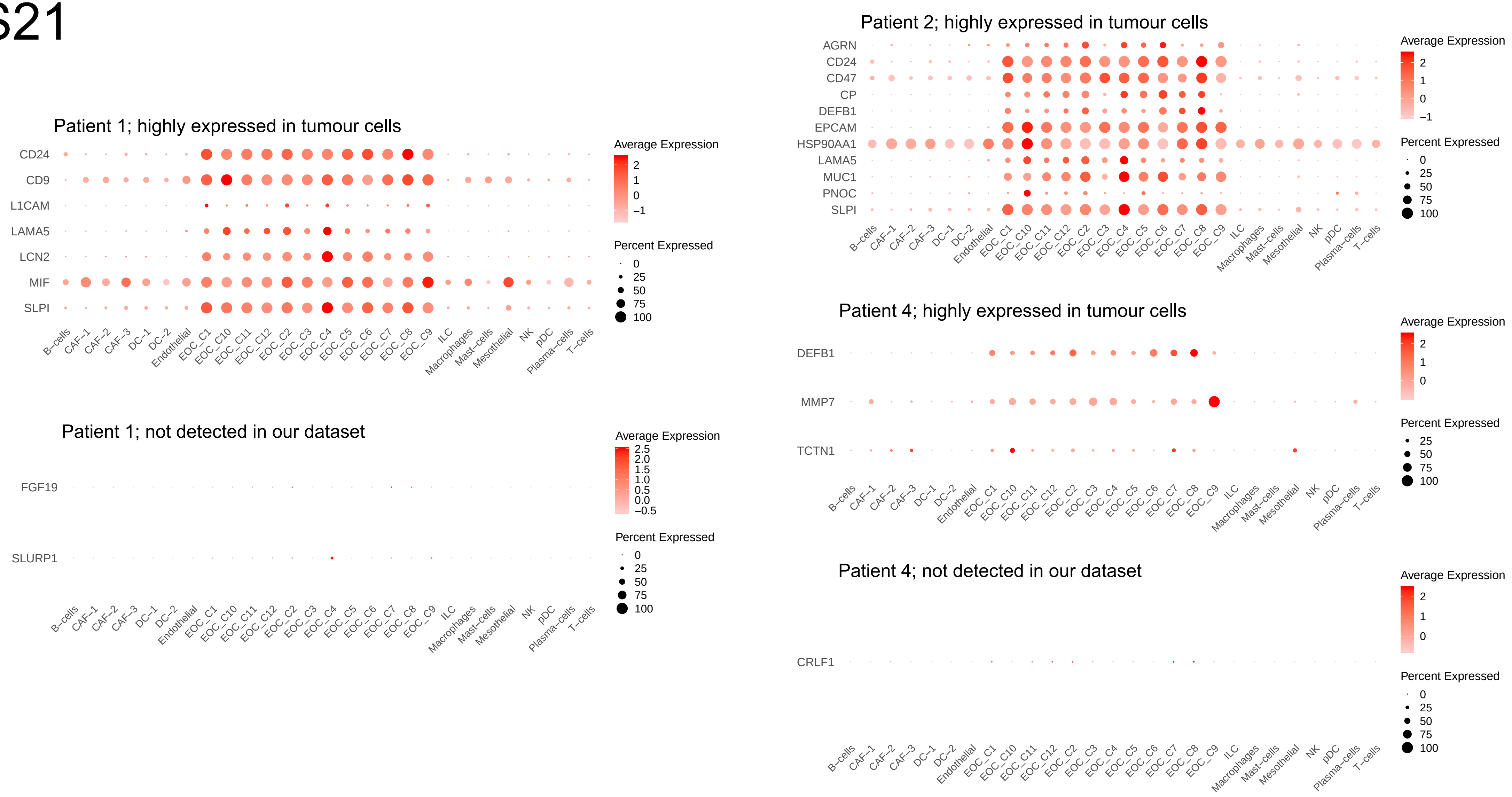


**Expression and detection rates of putative tumour intrinsic ligands in Zhang *et al.* scRNA-seq dataset.** Ligands identified in different samples are shown separately and are further split into ligands that were most highly expressed in tumour cells in our scRNA-seq dataset and ligand that were deemed plausibly subclone-specific as they were not detected in any of the cell types identified in our scRNA-seq dataset. Note that the latter did not show strong expression in any of the non-malignant subtypes. EOC, epithelial ovarian carcinoma.

### Fig. S21 continued

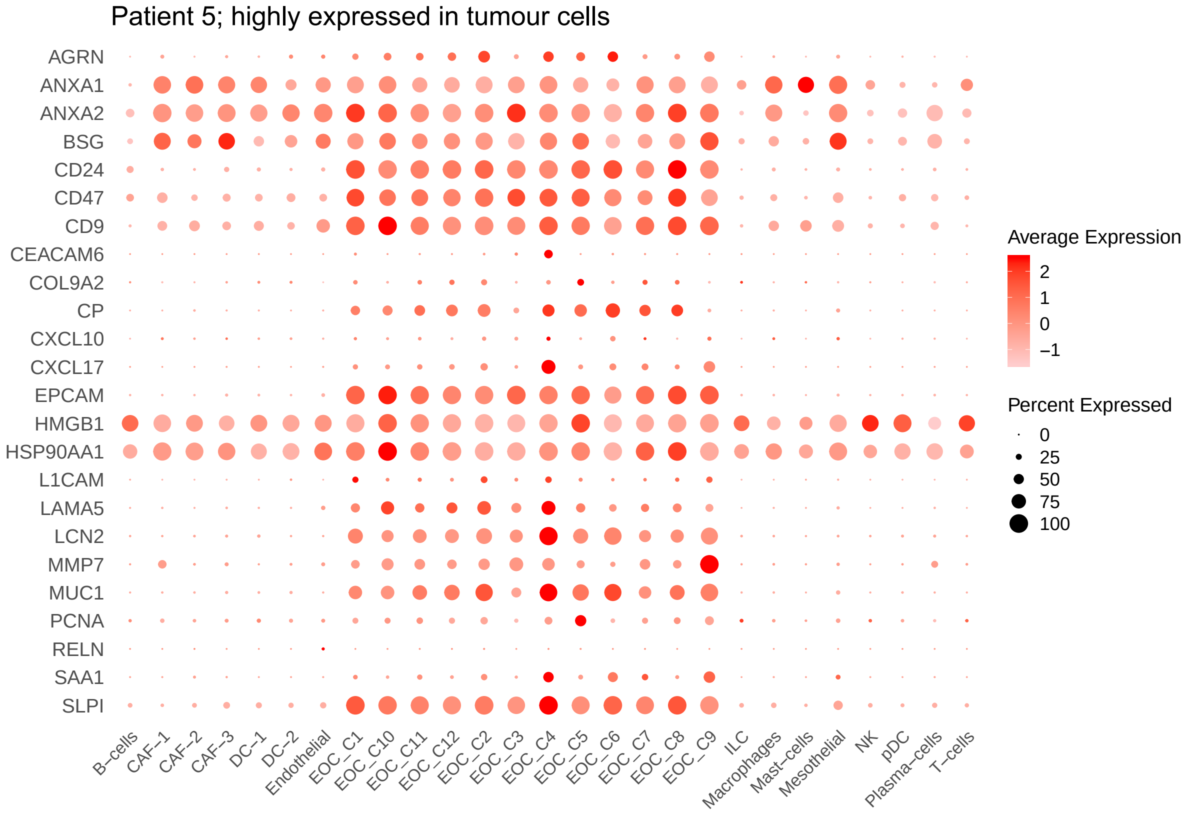


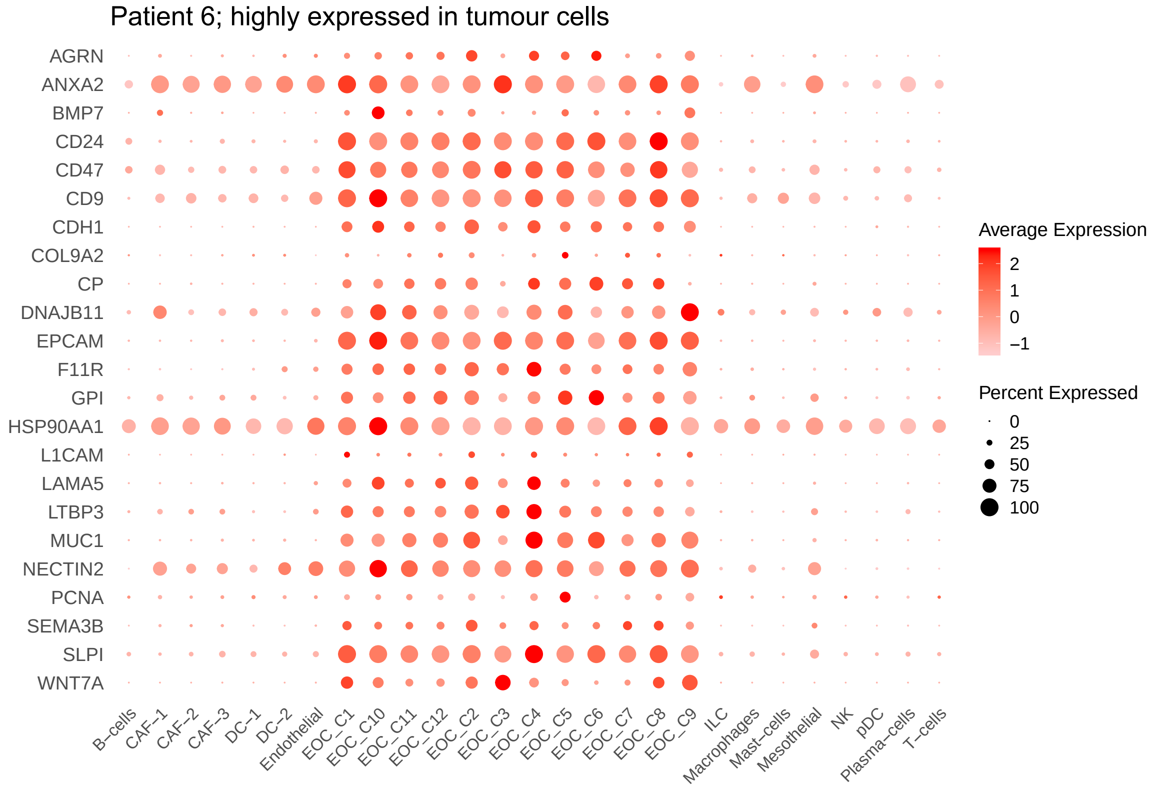


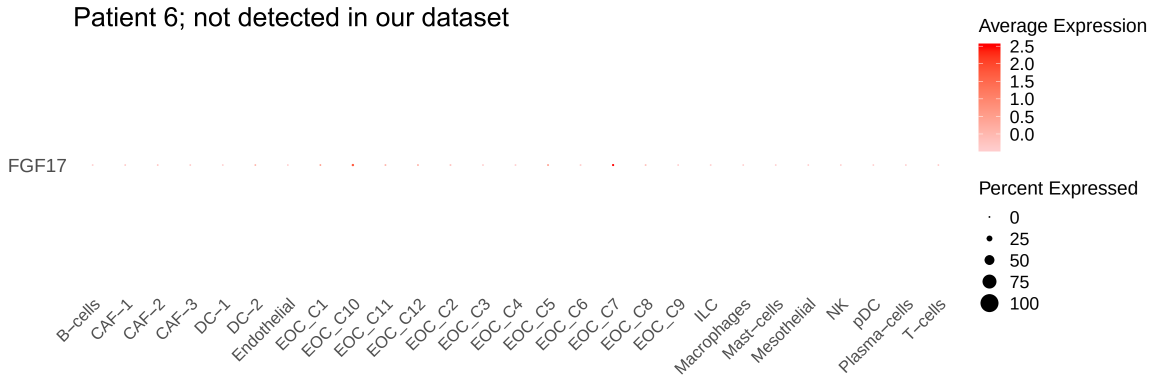

### Fig. S22

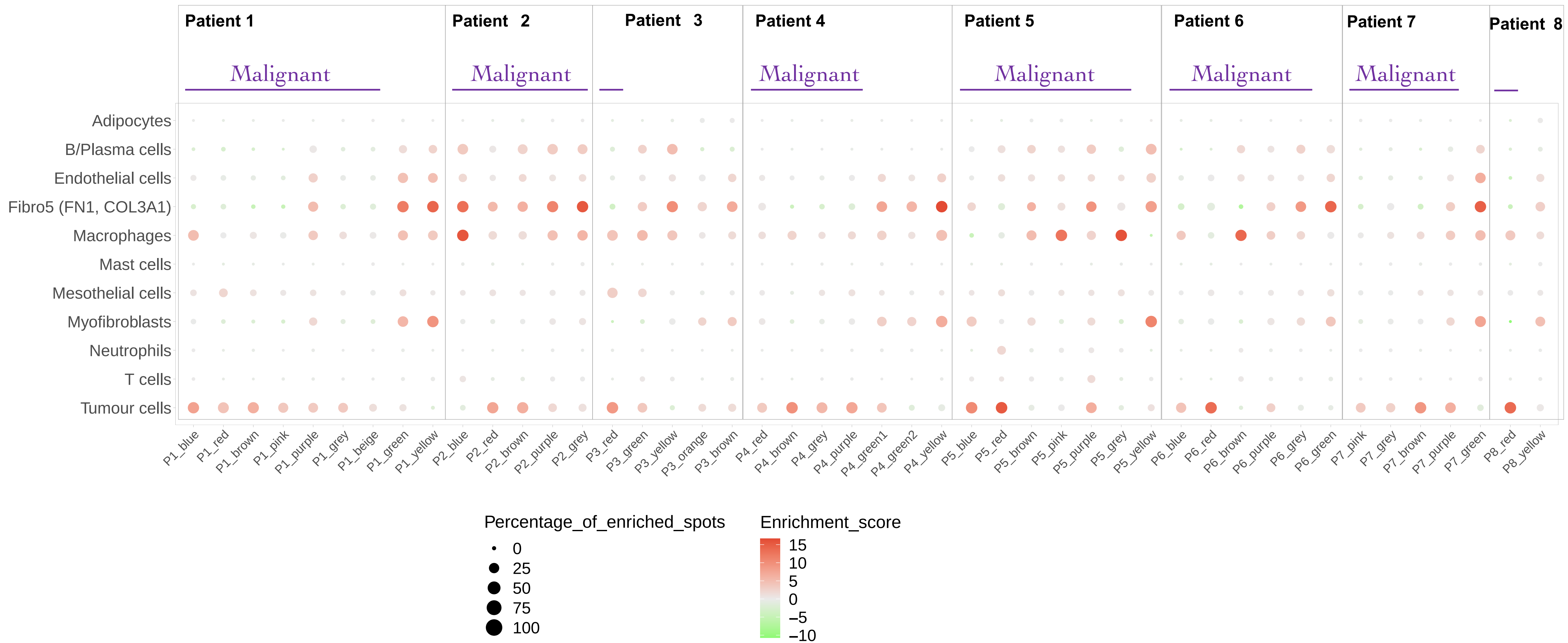


**Giotto cell type enrichment in inferCNV clusters.** Dot colour shows PAGE enrichment scores averaged across all spots in each inferCNV cluster and each cell type; dot size shows percentage of spots in each inferCNV cluster that were significantly enriched for the corresponding cell type (significance determined using a threshold of 0.05 on Benjamini-Hochberg adjusted Giotto p-values). All malignant and non-malignant inferCNV clusters are shown for each sample, purple bars indicate the malignant clusters.

### Fig. S23

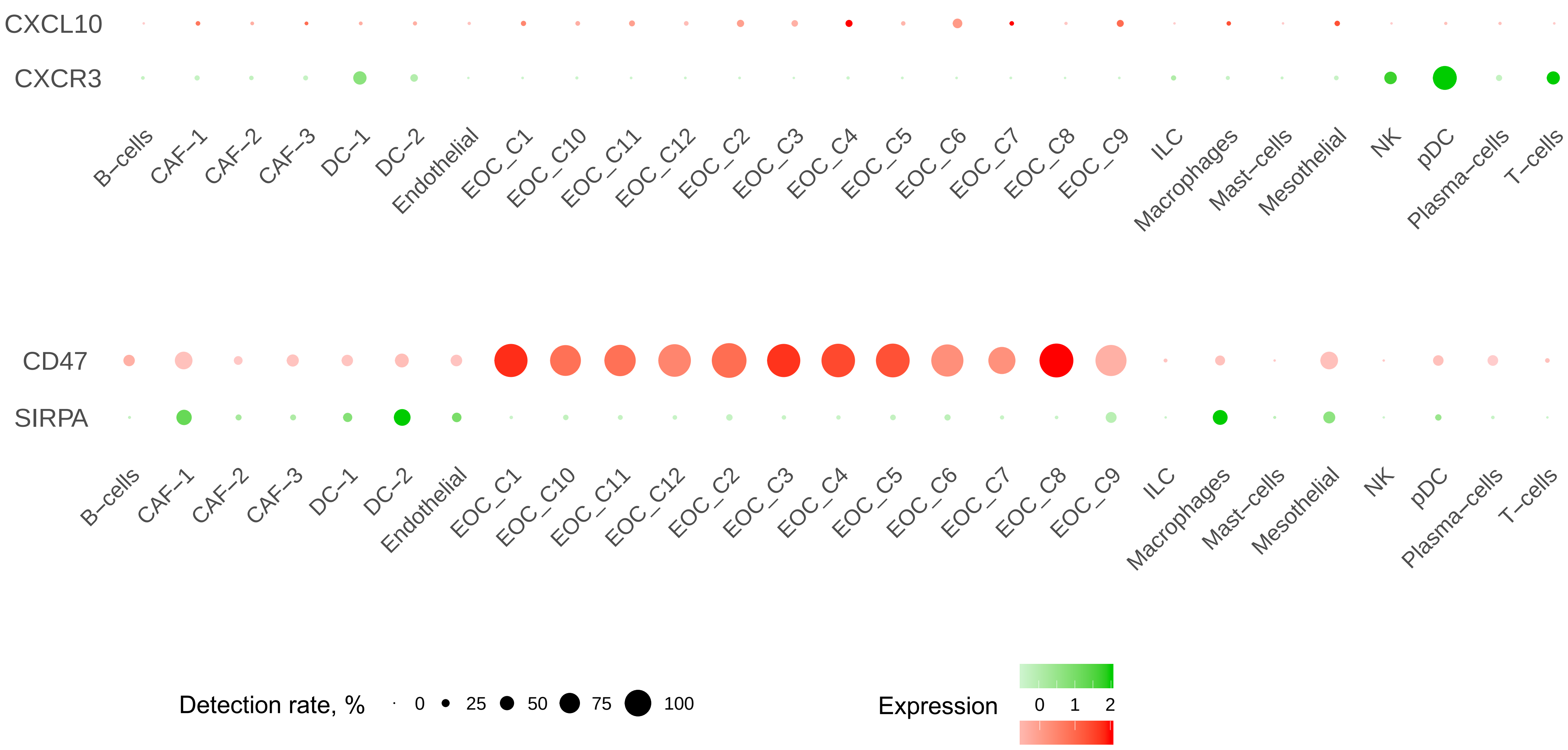


**Expression and detection rates of *CXCL10-CXCR3* and *CD47-SIRPA* ligand-receptor pairs in Zhang *et al.* scRNA-seq dataset.**

### Fig. S24

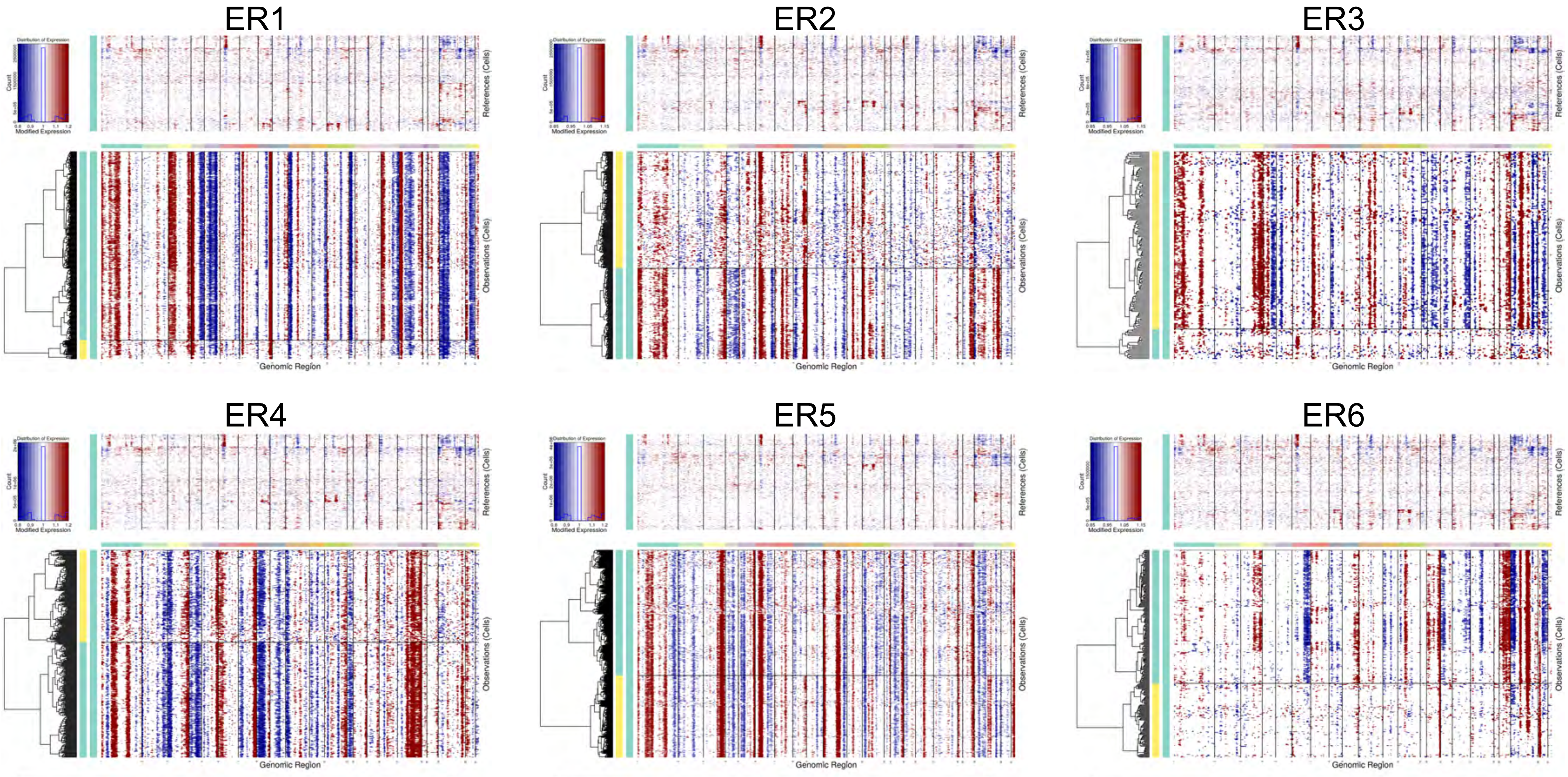


**InferCNV heatmaps generated for Visium samples from Stur *et al*. study.**

### Fig. S24 continued

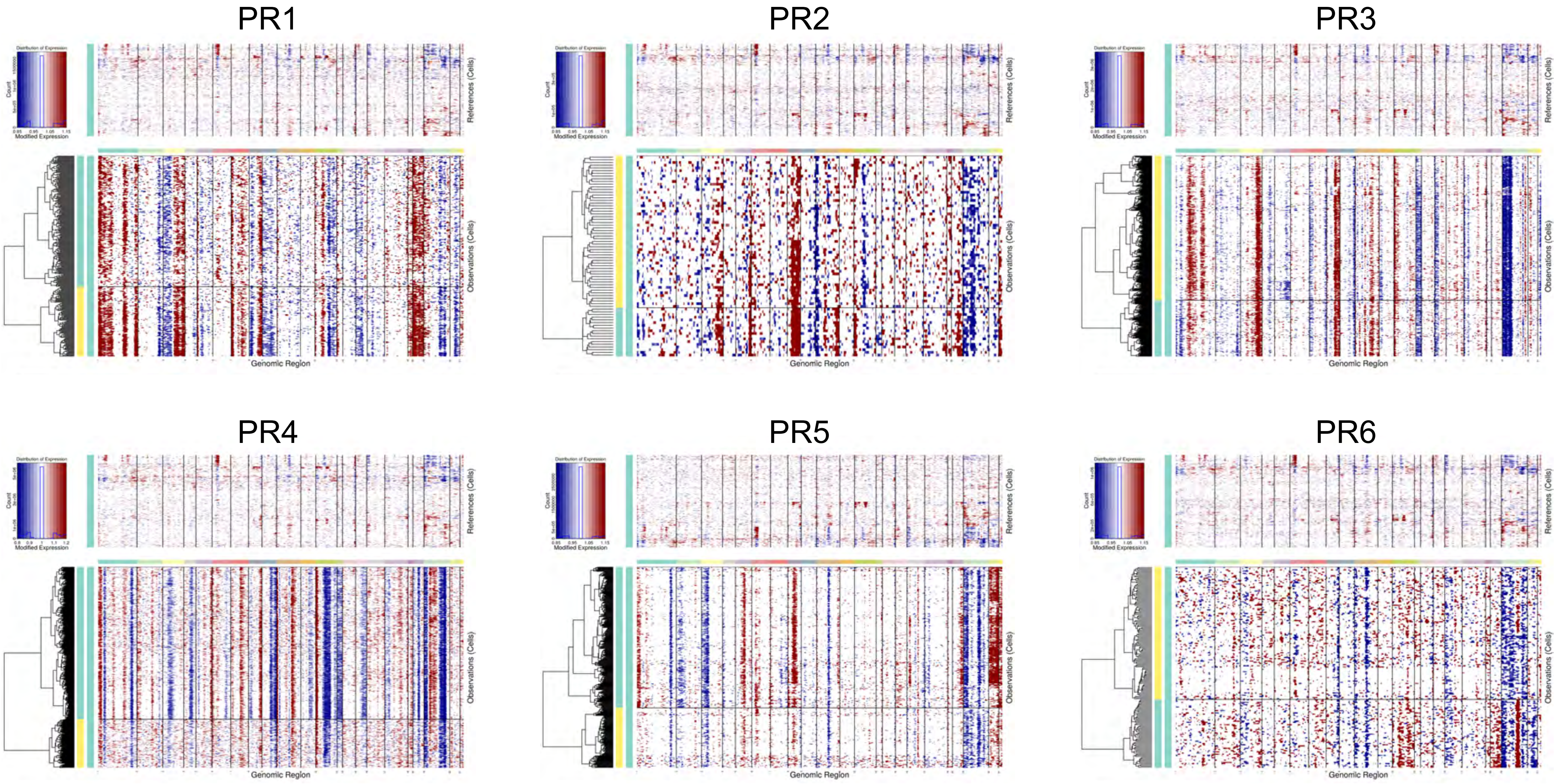

### Fig. S25

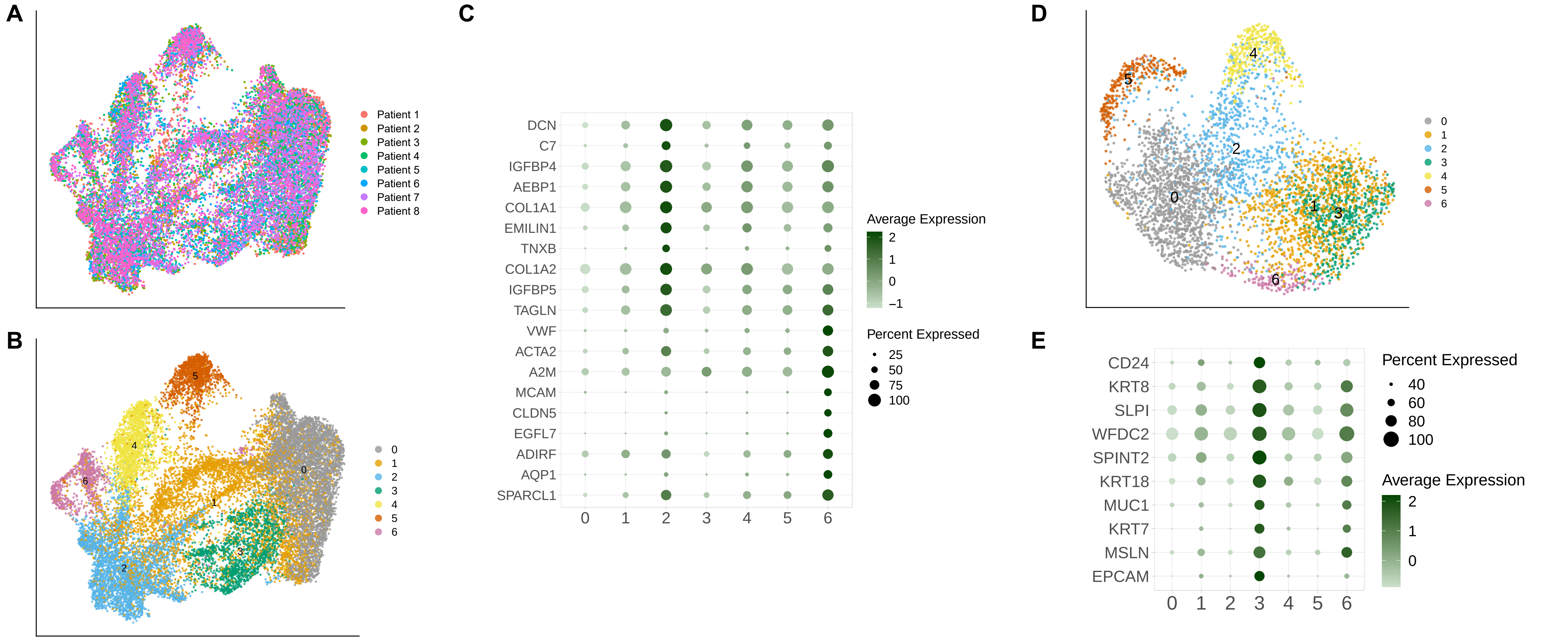


**Integrated gene expression analysis of eight Visium samples. a)** UMAP of eight integrated samples coloured by the sample of origin. **b)** UMAP of eight integrated samples coloured by the cluster. **c)** Top overexpressed genes in clusters 2 and 6 (candidate stromal dataset). **d)** Subclustering of candidate stromal dataset (clusters 2 and 6 from (**b**). **e)** Top overexpressed genes in cluster 3 of the candidate stromal dataset.

### Fig. S26

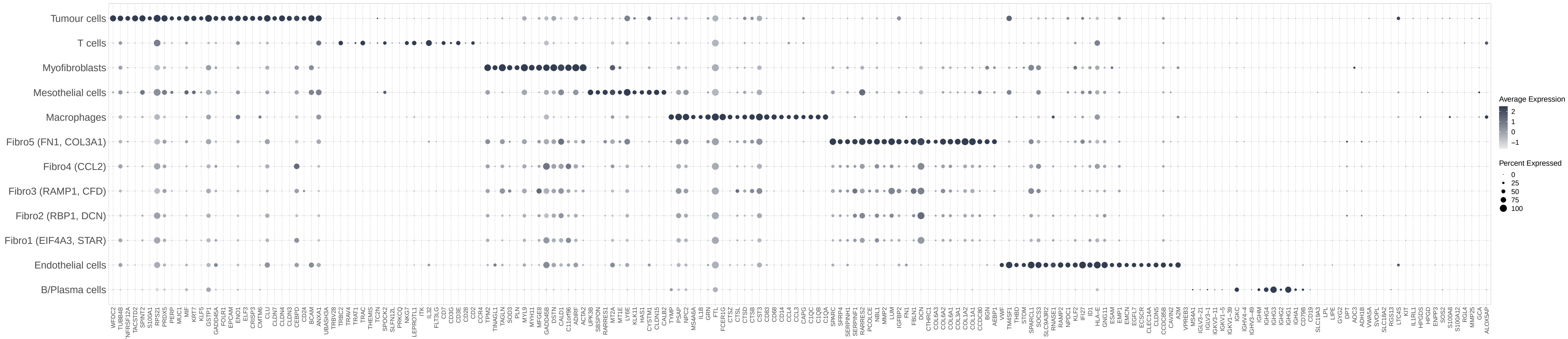


**Expression of marker genes used for Giotto cell type enrichment analysis in the scRNA-seq dataset.**
